# A minimal kinase–phosphatase system and its lipid substrates self-organize into dynamic patterns

**DOI:** 10.64898/2026.07.27.740984

**Authors:** Ting-Sung Hsieh, Beatrice Ramm, Qiwei Yu, Jonathon L. Yuly, James P. Pfister, Yinhao Wu, Junzhou Huang, Yigal Meir, Ned S. Wingreen, Vincent S. Tagliabracci

## Abstract

Competing lipid kinases and phosphatases are critical for organizing cellular membranes, but whether a minimal system can autonomously organize proteins and lipids into dynamic spatiotemporal patterns is unknown. Here, we report the in vitro reconstitution of the *Legionella* phosphatidylinositol (PI) 3-kinase MavQ and PI 3-phosphatase SidP. Together with their lipid substrates, PI and PI 3-phosphate (PI3P), these enzymes form a minimal self-organizing system that generates ATP-dependent spatiotemporal patterns, including traveling waves, on model membranes. These behaviors arise from MavQ’s cooperative membrane binding, SidP’s phosphatase activity, and the continual interconversion and redistribution of PI and PI3P within a conserved membrane pool. A reaction–diffusion model reproduces the observed dynamics and predicts that lipid conservation prevents patterns from propagating across membrane discontinuities, which we verify experimentally. Together, these findings establish enzymatic modification of membrane lipids as a distinct molecular strategy for biological pattern formation.

## Main Text

Dynamic spatial patterns of proteins and lipids underlie many cellular processes, including cell division, cell polarity, and cell migration (*1, 2*). These patterns can arise through self-organization, a fundamental principle by which order emerges from local molecular interactions in complex chemical and biological systems (*3*). Self-organization can be achieved through reaction–diffusion mechanisms, in which diffusing and interacting chemical species spontaneously generate spatiotemporal patterns from an initially homogeneous state (*4*). Identifying the minimal molecular components sufficient for reaction–diffusion-driven pattern formation is essential for understanding the physical principles that govern cellular organization.

In vitro reconstitution provides a rigorous approach to identifying minimal molecular components and dissecting the underlying mechanisms. To date, only a handful of intracellular pattern-forming systems have been reconstituted from purified components, including the MinDE system from *Escherichia coli* (*5*), one of the best-characterized examples of dynamic intracellular pattern formation. MinD and MinE oscillate between the cell poles to position the division site (*6*) and form traveling waves on model membranes (*7*). Only four components— MinD, MinE, adenosine triphosphate (ATP), and a charged membrane—are sufficient to reconstitute pattern formation in vitro (*7*), transforming the MinDE system into a tractable model for mechanistic studies (*5*). The success of the MinDE system highlights the power of this reductionist approach but extending it to more complex cellular systems is far from straightforward. For example, in eukaryotic cells, spatiotemporal patterns typically emerge from the coordinated activities of numerous interacting components, where extensive molecular redundancy makes it difficult to define the minimal components sufficient for pattern formation in vivo.

Phosphoinositide signaling provides a compelling example of this complexity. Lipid kinases and phosphatases convert phosphatidylinositol (PI) into various phosphoinositide species, which exhibit distinct, dynamic membrane localizations crucial for defining compartmental identity and regulating membrane trafficking in eukaryotic cells (*8*). Evidence indicates that phosphoinositide kinases and phosphatases drive the dynamic patterning of phosphoinositides in cellular membranes (*9–11*), a process theoretically predicted to arise via reaction–diffusion mechanisms (*2*). However, because phosphoinositide signaling involves complex regulatory networks (*12*), determining whether a minimal kinase–phosphatase pair is sufficient to generate dynamic patterns has remained challenging. Indeed, while the in vitro reconstitution of a competitive eukaryotic lipid kinase–phosphatase system successfully demonstrated spatial composition patterns, these were restricted to stationary domains arising from intrinsic bistability and stochastic symmetry breaking (*13*). Whether a kinase–phosphatase pair can autonomously sustain dynamic spatiotemporal patterns on a membrane remains an open question.

Intracellular bacterial pathogens provide an attractive system for identifying minimal self-organizing modules because they have evolved to manipulate host signaling pathways using relatively few effector proteins. We previously identified one such system in the *Legionella* effector repertoire consisting of a two-component module that self-organizes to remodel the host endoplasmic reticulum (ER). In the absence of other bacterial effector proteins, the *Legionella* PI 3-kinase MavQ and PI 3-phosphatase SidP generate spatiotemporal patterns and exhibit oscillatory behavior on the ER membrane (**Fig. S1A and movie S1**) while simultaneously driving rapid turnover of PI 3-phosphate (PI3P) **(Fig. S1B and movie S2**) (*14*). These observations suggest that the MavQ–SidP system and its lipid substrates, PI and PI3P, constitute a strong candidate for a minimal, self-organizing reaction–diffusion system. However, whether these components are sufficient to generate dynamic patterns in the absence of additional host factors remained unknown. Here, we report the in vitro reconstitution of the MavQ–SidP system on model membranes. By combining quantitative experiments with a reaction–diffusion model, we show that this kinase–phosphatase pair, together with its lipid substrates PI and PI3P, is sufficient to drive autonomous dynamic patterns, providing insight into how competing lipid-modifying enzymes can direct self-organization on membranes.

### The *Legionella* PI 3-kinase–phosphatase system MavQ–SidP self-organizes and forms spatiotemporal patterns on membranes

To investigate pattern formation by MavQ and SidP, we used an in vitro reconstitution assay where a supported lipid bilayer (SLB) was formed on a coverslip in an open reaction chamber filled with buffer solution (**Fig. 1A**). In assays containing PI in the SLB, ATP addition caused MavQ to bind the membrane and gradually form a uniform protein layer (**Fig. S2, A and B**), whereas assays containing both MavQ and SidP underwent symmetry breaking and generated dynamic patterns (**Fig. 1B and movie S3**). Conversely, in the absence of any of the four components—MavQ, SidP, PI, or ATP—pattern formation was abolished (**Fig. S2, C and D, and movie S3**). Following ATP addition, pattern formation began with the emergence of dense, motile MavQ patches with internal density gradients (**Fig. 1, B and C, and movie S3**). Over time, these patches self-organized into traveling waves (**Fig. 1, B and D, Fig. S2E, and movie S3**) that, under most conditions, ultimately condensed into quasi-stationary patterns (**Fig. 1B and movie S3**), including spots and labyrinths (**Fig. 1E**). The formation of these quasi-stationary patterns was not an artifact of phototoxicity during time-lapse imaging (**Fig. S2F and movie S4**). Using total internal reflection fluorescence microscopy, we confirmed that SidP colocalized and traveled with MavQ within these dynamic patterns (**Fig. 1F and movie S5**), consistent with their behavior when expressed in eukaryotic cells (**Fig. S1A and movie S1**) (*14*). Thus, MavQ, SidP, PI, and ATP are sufficient for dynamic, self-organized pattern formation.

**Fig. 1.**
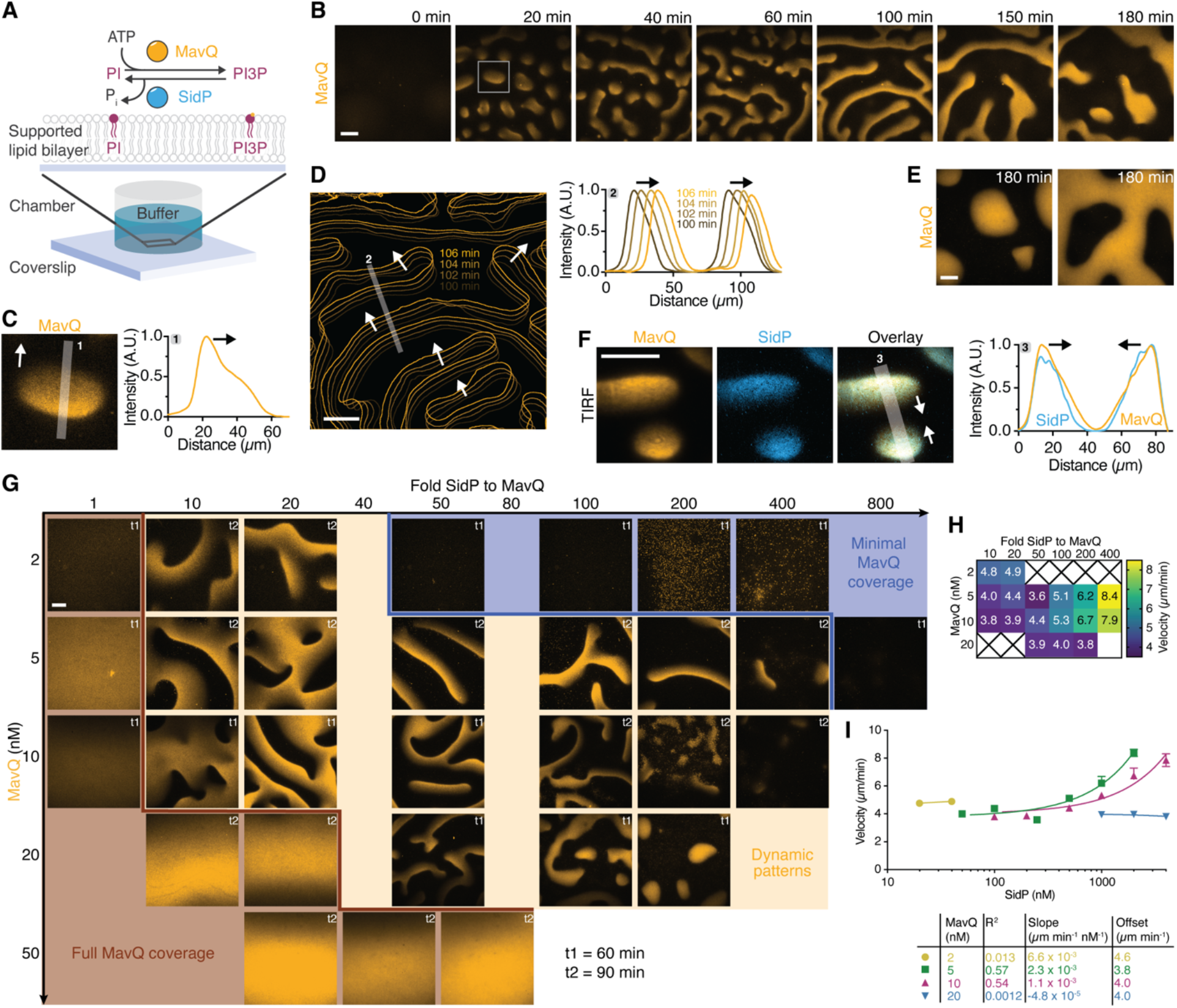
The *Legionella* PI 3-kinase–phosphatase system MavQ–SidP self-organizes and forms spatiotemporal patterns on membranes. **(A)** Schematic representation of the in vitro reconstitution assay. **(B)** Time-lapse confocal images showing the emergence of dynamic patterns and waves following ATP addition immediately before 0 min. See also the first panel of movie S3. **(C)** Close-up of the boxed area in (B) at 20 min showing a MavQ patch with an internal density gradient, which is visualized by the corresponding intensity profile (taken along the gray line 1; smoothed). Arrows indicate the direction of patch movement. **(D)** Color-coded overlay of pattern contours and corresponding intensity profiles (taken along the gray line 2; smoothed) from four consecutive time points starting at 100 min in (B), showing wave propagation. **(E)** Confocal images of quasi-stationary MavQ patterns observed 180 min after ATP addition, showing spots (left panel) and labyrinths (right panel). **(F)** Total internal reflection fluorescence images and corresponding intensity profile (taken along the gray line 3; smoothed) showing colocalization of MavQ and SidP in the patterns. Arrows indicate the direction of wave propagation. See also movie S5. **(G)** Confocal images from two representative time points illustrating the three patterning regimes as a function of MavQ concentration and the SidP-to-MavQ molar ratio. Brightness and contrast are comparable within each row. See also Fig. S3A and movie S6. **(H)** Heatmap showing wave velocity as a function of MavQ concentration and the SidP-to-MavQ molar ratio. Crosses (×) indicate conditions under which no dynamics were observed. Data points: mean from at least 6 tracked waves per condition across 4 independent experiments. **(I)** Wave velocity as a function of SidP concentrations at 2, 5, 10, and 20 nM MavQ. Data points and bars: mean and SEM from at least 6 tracked waves per condition across 4 independent experiments; curves: linear regression with parameters indicated in the table below. Initial lipid composition: 95% DOPC, 5% 18:1 PI, and an extra 0.01% ATTO 647N PE. Proteins: 10 nM MavQ with 1 µM SidP (B to E), 10 nM MavQ with 2 µM SidP (F), or as indicated (G to I). For all panels, MavQ was doped with 25% mEGFP-MavQ. For panel (F), SidP was doped with 25% Alexa Fluor 568-labeled SidP. ATP was added immediately before the start of image acquisition. Scale bars: 50 µm.

We systematically varied the bulk concentrations of MavQ and SidP to map how these two parameters shape the resulting spatiotemporal patterns. This survey revealed distinct regimes, defined as ranges of MavQ concentration and SidP-to-MavQ molar ratio that gave rise to qualitatively different patterns (**Fig. 1G, Fig. S3A, and movie S6**). At the extremes, two limiting behaviors emerged: at low SidP-to-MavQ ratios or high MavQ concentrations, MavQ uniformly coated the membrane, occasionally following a transient period of pattern formation. Conversely, high SidP-to-MavQ ratios and low MavQ concentrations resulted in minimal membrane coverage. Consistent with these observations, increasing MavQ concentrations increased MavQ intensities on membranes, whereas increasing the SidP-to-MavQ ratio had the opposite effect (**Fig. S3B**). Between these extremes, we observed a broad pattern-forming regime characterized by dynamic patterns and a high coefficient of variation in MavQ intensity across the field of view (**Fig. S3C**). Within this pattern-forming regime, wavefronts propagated at velocities ranging from 3.6 to 8.4 μm/min (**Fig. 1H**). At MavQ concentrations of 5 nM or 10 nM, raising the SidP concentration increased wave velocity (**Fig. 1I**). Wave velocity reached a maximum near the boundary of the minimal-MavQ-coverage regime, where periodic wave trains transitioned into sporadic wavefronts (**Fig. 1G and movie S6**).

A key distinction between the MavQ–SidP and MinDE systems is the role of the membrane. Whereas the membrane serves primarily as a passive platform in the MinDE system, the MavQ– SidP system actively interconverts its lipid substrates. To examine how lipid substrate availability influences pattern formation, we varied both the PI concentration in the membrane and the MavQ concentration in solution while maintaining a constant SidP-to-MavQ ratio. Increasing either the PI concentration in the membrane or the MavQ concentration in solution increased both MavQ membrane recruitment and membrane coverage (**Fig. S4, A and B, and movie S7**). At low PI concentrations (≤ 1%), we frequently observed spots and stripes with wavy edges (**Fig. S4C and movie S7**), reminiscent of the capillary waves predicted to occur on active condensates (*15*). Collectively, these results establish that the reconstituted MavQ–SidP system self-organizes into diverse patterns whose emergence and dynamics are controlled by both protein and lipid substrate abundance.

### MavQ binds PI3P-positive membranes cooperatively via CTD-mediated interactions

We hypothesized that MavQ’s membrane binding is cooperative, a potential source of the nonlinear kinetics required for reaction–diffusion-driven pattern formation of the MavQ–SidP system (*4, 5, 16*). To test this idea, we measured MavQ membrane binding on silica bead-supported lipid bilayers (BSLBs) and determined cooperativity from saturation binding curves (*17*). In the presence of ATP, MavQ produced PI3P on PI-containing BSLBs (**Fig. 2A and movie S8**) and bound cooperatively to the membrane, with a Hill coefficient (*n*_H_) greater than 1 (**Fig. 2B**). In contrast, in the presence of the non-hydrolyzable ATP analog AMPPCP (adenosine-5’-[(β,γ)-methyleno]triphosphate), MavQ neither generated PI3P nor bound to PI-containing BSLBs (**Fig. S5A and movie S9**), suggesting that stable membrane association requires either ATP hydrolysis or PI3P generated by MavQ.

**Fig. 2.**
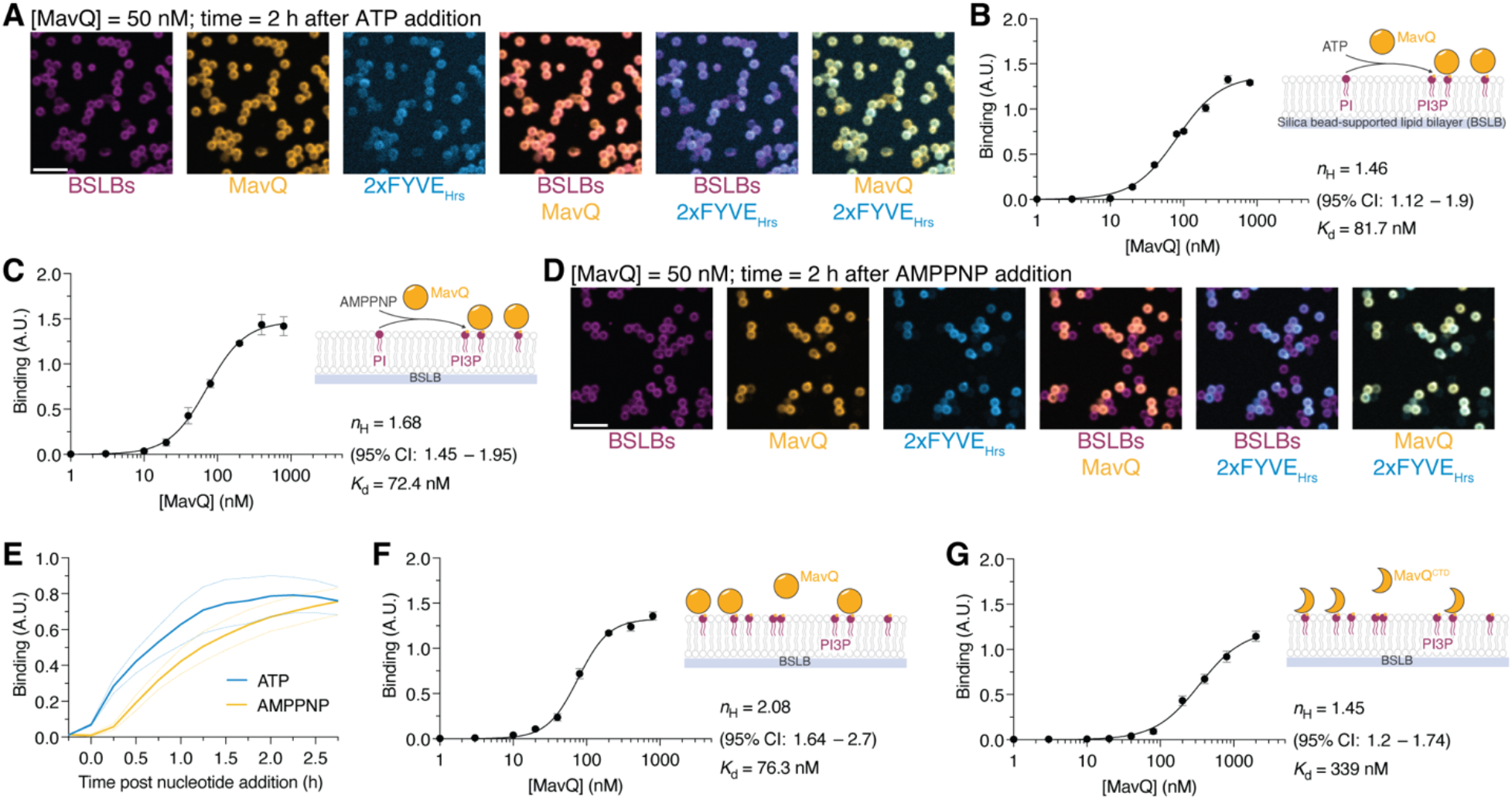
MavQ binds PI3P-positive membranes cooperatively via CTD-mediated interactions. **(A)** Confocal images of MavQ membrane binding and PI3P production on PI-containing bead-supported lipid bilayers (BSLBs), taken 2 h after ATP addition. PI3P was visualized with the probe mTagBFP2-2xFYVE_Hrs_. See also movie S8. (**B** and **C**) Quantification of MavQ binding at equilibrium to PI-containing BSLBs in the presence of ATP (B) or AMPPNP (C). Each data point represents the mean fluorescence intensity of at least 600 beads. Data points and bars: mean and SEM of 4 independent experiments; curves: nonlinear regression using a specific binding model with Hill slope. (**D**) Confocal images of MavQ membrane binding and PI3P production on PI-containing BSLBs, taken 2 h after AMPPNP addition. PI3P was visualized using the probe mTagBFP2-2xFYVE_Hrs_. See also movie S10. (**E**) Time course of MavQ binding to BSLBs. Nucleotides were added immediately before time zero. Curves: mean and SEM of 3 independent experiments. (**F** and **G**) Quantification of MavQ (F) or MavQ^CTD^ (G) binding at equilibrium to PI3P-containing BSLBs. Each data point represents the mean fluorescence intensity of at least 600 beads. Data points and bars: mean and SEM of 4 independent experiments; curves: nonlinear regression using a specific binding model with Hill slope. Initial lipid compositions: 95% DOPC, 5% 18:1 PI (A to E) or 18:1 PI3P (F and G), and an extra 0.01% ATTO 647N PE. Proteins: 50 nM MavQ, doped with 25% mEGFP-MavQ, and 50 nM mTagBFP2-2xFYVE_Hrs_ (A, D, and E); MavQ, doped with 25% mEGFP-MavQ, or MavQ^CTD^, doped with 25% mEGFP-MavQ^CTD^, at several concentrations through saturation (B, C, F, and G). Scale bars: 20 µm.

Notably, MavQ membrane binding remained cooperative in the presence of the slowly hydrolyzable ATP analog AMPPNP (adenosine-5’-[(β,γ)-imido]triphosphate) (**Fig. 2C**), but the binding was slower and occurred in an all-or-none manner compared with binding in the presence of ATP (**Fig. 2, D and E, and movie S10**). We propose that this behavior arises because slow AMPPNP hydrolysis likely makes PI3P production stochastic. Under these conditions, the rare production of the first few PI3P molecules on an individual silica bead may nucleate rapid cooperative recruitment of additional MavQ molecules, giving rise to the observed all-or-none binding.

We next asked whether the observed cooperativity could be explained by PI3P production– binding feedback (*14*), whereby MavQ recruitment is enhanced by its own product, PI3P, through its PI3P-binding C-terminal domain (CTD, residues 581 to 853). However, our results indicate that this feedback loop is not required for cooperative membrane binding. First, in the absence of ATP, MavQ membrane binding remained cooperative on BSLBs that already contained PI3P (**Fig. 2F**), a condition that decouples PI3P production from membrane binding. Second, the isolated CTD of MavQ, which lacks the kinase domain, was sufficient to bind cooperatively to PI3P-containing BSLBs (**Fig. 2G**). Therefore, MavQ-dependent PI3P production initiates and facilitates MavQ membrane recruitment, whereas cooperativity itself is an intrinsic property of CTD-mediated interactions between MavQ molecules.

### SidP’s PI 3-phosphatase activity provides the negative feedback required for pattern formation

Reaction–diffusion-driven pattern formation also requires negative feedback (*18*). We previously found that both SidP’s PI 3-phosphatase activity and its non-catalytic MavQ-interacting CTD (SidP^CTD^, residues 664 to 822) reduce MavQ association with the ER (*14*), so either mechanism might provide the negative feedback required for pattern formation. To identify the source of the negative feedback, we tested catalytically inactive SidP^C554S^, which, unlike wild-type SidP, failed to drive spatiotemporal patterning of MavQ on SLBs (**Fig. S5B**).

Unexpectedly, however, SidP^C554S^, which retains the MavQ-interacting CTD, enhanced MavQ membrane binding, resulting in faster binding kinetics and a higher, biphasic maximal binding capacity (**Fig. S5C**). This effect was specific: the non-interacting control protein, Protein A, did not similarly enhance MavQ binding (**Fig. S5C**). Moreover, the isolated SidP^CTD^ was sufficient to enhance MavQ membrane binding (**Fig. S5C**). Similar results were obtained using BSLBs (**Fig. S5D**), where MavQ achieved binding equilibrium more rapidly than on flat SLBs. Because SidP directly interacts with both MavQ (*14, 19*) and PI3P (**Fig. S5E**), these results suggest that SidP promotes multivalent, high-avidity membrane association of MavQ by simultaneously engaging both the protein and the lipid. Thus, although the SidP^CTD^ enhances MavQ membrane binding, SidP phosphatase activity provides the negative feedback required for pattern formation.

### Self-organization of MavQ and SidP drives spatiotemporal patterning of phospholipids in membranes

We previously observed that PI3P lipid waves accompany MavQ and SidP protein waves on the ER in cells (**Fig. S1B and movie S2**) (*14*). We therefore asked whether PI3P is similarly patterned by the reconstituted MavQ–SidP system in vitro. Using the fluorescent PI3P probe 2xFYVE_Hrs_ (**Fig. 3A**), we found that PI3P patterns colocalized with MavQ patterns, and PI3P lipid waves propagated in step with the MavQ protein waves (**Fig. 3B and movie S11**). This dynamic colocalization could arise from two non-mutually exclusive mechanisms: localized PI3P synthesis and turnover by the opposing kinase and phosphatase activities of MavQ and SidP, or active concentration and cotransport of PI3P through the PI3P-binding domains of the proteins.

**Fig. 3.**
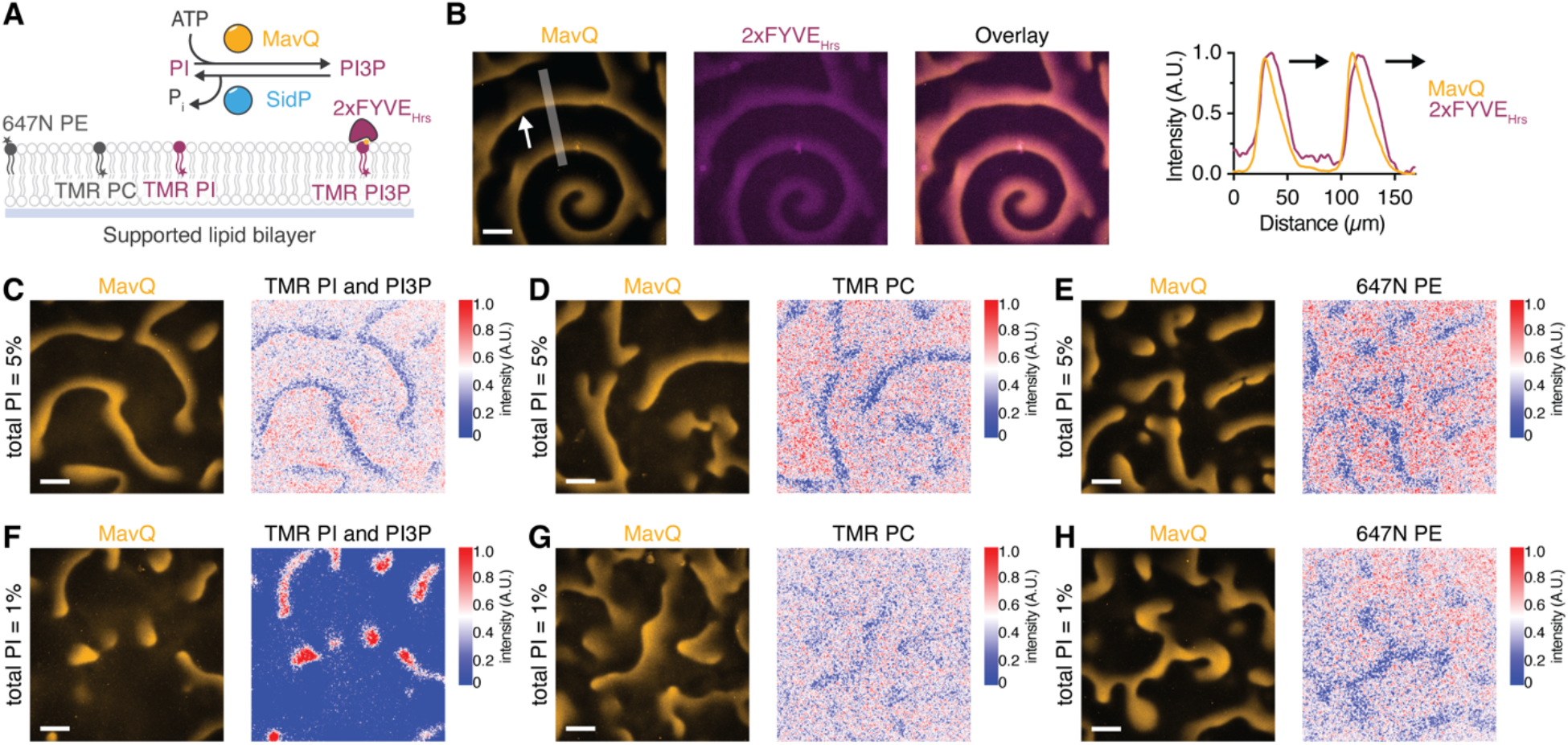
Self-organization of MavQ and SidP drives spatiotemporal patterning of phospholipids in membranes. (**A**) Schematic representation of the in vitro reconstitution assay for tracking phospholipid dynamics during MavQ–SidP pattern formation. TopFluor TMR PC, PI, or PI3P (TMR PC, PI, or PI3P); ATTO 647N PE (647N PE). (**B**) Confocal images and corresponding intensity profile (taken along the gray line; smoothed) showing colocalization of MavQ and PI3P within patterns. PI3P was visualized with the probe mTagBFP2-2xFYVE_Hrs_. Arrows indicate the direction of wave propagation. See also movie S11. (**C** to **E**) Confocal images and heat maps of MavQ patterns and the underlying phospholipid distribution in membranes containing 5% total PI. The combined PI and PI3P pool was tracked using TMR PI (C), while TMR PC and 647N PE served as non-substrate lipid controls for lipid sorting (D and E). See also movies S12, S14, and S15. (**F** to **H**) Confocal images and heat maps of MavQ patterns and phospholipid distribution in membranes containing 1% total PI. The combined PI and PI3P pool was tracked using TMR PI (F), while TMR PC and 647N PE served as non-substrate lipid controls for lipid sorting (G and H). See also movies S16, S18, and S19. Initial lipid composition: 95% DOPC, 5% 18:1 PI, and an extra 0.01% ATTO 647N PE (B and E); 95% DOPC, 4.9% 18:1 PI, and 0.1% TopFluor TMR PI (C); 94.9% DOPC, 5% 18:1 PI, and 0.1% TopFluor TMR PC (D); 99% DOPC, 0.9% 18:1 PI, and 0.1% TopFluor TMR PI (F); 98.9% DOPC, 1% 18:1 PI, and 0.1% TopFluor TMR PC (G); 99% DOPC, 1% 18:1 PI, and an extra 0.01% ATTO 647N PE (H). Proteins: 20 nM MavQ and 2 µM SidP (B to H), with the addition of 20 nM mTagBFP2-2xFYVE_Hrs_ for (B). For all experiments, MavQ was doped with 25% mEGFP-MavQ, and ATP was added immediately before the start of image acquisition. Scale bars: 50 µm.

To determine whether MavQ and SidP transport their lipid substrates, we incorporated a small fraction of fatty-acid-tail-labeled TopFluor TMR PI, which retains an unmodified inositol headgroup and can therefore be phosphorylated, allowing the collective visualization of PI and PI3P dynamics (**Fig. 3A**). At a total PI concentration of 5%, this fluorescent lipid, which reports the combined PI and PI3P pool, was weakly excluded from MavQ-dense regions (**Fig. 3C and movie S12**), even though the PI3P probe 2xFYVE_Hrs_ remained enriched there (**Fig. S6A and movie S13**). This exclusion was not substrate specific, as the non-substrate lipids TopFluor TMR phosphatidylcholine (PC) and ATTO 647N phosphatidylethanolamine (PE) displayed similar behavior (**Fig. 3, D and E, and movies S14 and S15)**. We attribute this effect to steric repulsion from dense protein layers or diffusiophoretic transport driven by membrane protein gradients, phenomena previously observed in the MinDE system (*20–22*).

However, a different behavior emerged when the total PI concentration was reduced from 5% to 1%, approaching a 1:1 ratio with the total amount of MavQ in the assay. Under these conditions, a fraction of the TopFluor TMR PI colocalized with and tracked MavQ patterns (**Fig. 3F and movie S16**), where the PI3P probe also accumulated (**Fig. S6B and movie S17**). In contrast, the non-substrate lipids TopFluor TMR PC and ATTO 647N PE remained weakly excluded from MavQ-dense regions (**Fig. 3, G and H, and movies S18 and S19)**. Together, these results show that the MavQ–SidP system patterns membrane phospholipids through two distinct mechanisms: general, nonspecific exclusion of membrane lipids and active concentration and transport of lipid substrates that becomes apparent when the substrate is not present in excess. This extends MavQ–SidP self-organization beyond the proteins themselves to the phospholipids they act on.

### A reaction–diffusion model captures pattern dynamics and predicts geometry-dependent wave propagation

To understand how MavQ, SidP, and their lipid substrates generate spatiotemporal patterns, we developed a reaction–diffusion model. Unlike MavQ and SidP, which exchange continuously with the surrounding solution, PI and PI3P remain confined to the membrane, so the total membrane pool of these lipids within a connected membrane patch is conserved. How this lipid conservation influences pattern dynamics remains unknown.

The model describes membrane-bound MavQ (*m*), SidP (*p*), and a conserved lipid field (*ϕ*), representing the combined pool of PI and PI3P. MavQ binds cooperatively to the membrane in a lipid-dependent manner, whereas SidP promotes MavQ dissociation through its phosphatase activity. SidP is recruited by membrane-bound MavQ through either direct protein–protein interactions or MavQ-generated PI3P before dissociating at a constant rate. Both proteins diffuse on the membrane, while the lipid field undergoes biased diffusion, producing a net motion of lipid substrates toward MavQ-dense regions. The favorable interactions between MavQ and the lipid substrates, including lipid-facilitated MavQ binding and MavQ-directed lipid motion, are crucial for the emergence of dynamical patterns. Together, these interactions define a minimal activator–inhibitor model on a conserved substrate (**Supplementary Text and Table S1**).

Our model qualitatively reproduced the main features of experimentally observed pattern dynamics (**Fig. 4A and movie S20**). Specifically, the model captured three distinct regimes: full MavQ coverage at low SidP-to-MavQ ratios or high MavQ concentrations, minimal MavQ coverage at high SidP-to-MavQ ratios and low MavQ concentrations, and dynamical pattern formation under intermediate conditions (**Fig. 4, A and B, and movie S20**). Within the pattern-forming regime, MavQ-dense regions were enriched in both SidP and the lipid substrates (**Fig. 4C**). Increasing the concentration of lipid substrates increased MavQ membrane coverage and promoted pattern formation (**Fig. S7, A and B, and movie S21**), consistent with our experimental observations (**Fig. S4A**). Likewise, wave velocity increased toward the phase boundary between the pattern-forming and minimal-MavQ-coverage regimes (**movie S20**), matching the behavior observed experimentally (**Fig. 1H**). Finally, the model reproduced the capillary-like fluctuations at the edges of MavQ-dense regions (**Fig. S7C**), reminiscent of those observed experimentally (**Fig. S4C**). Thus, a minimal reaction–diffusion model captures the principal features of the experimentally observed pattern dynamics.

**Fig. 4.**
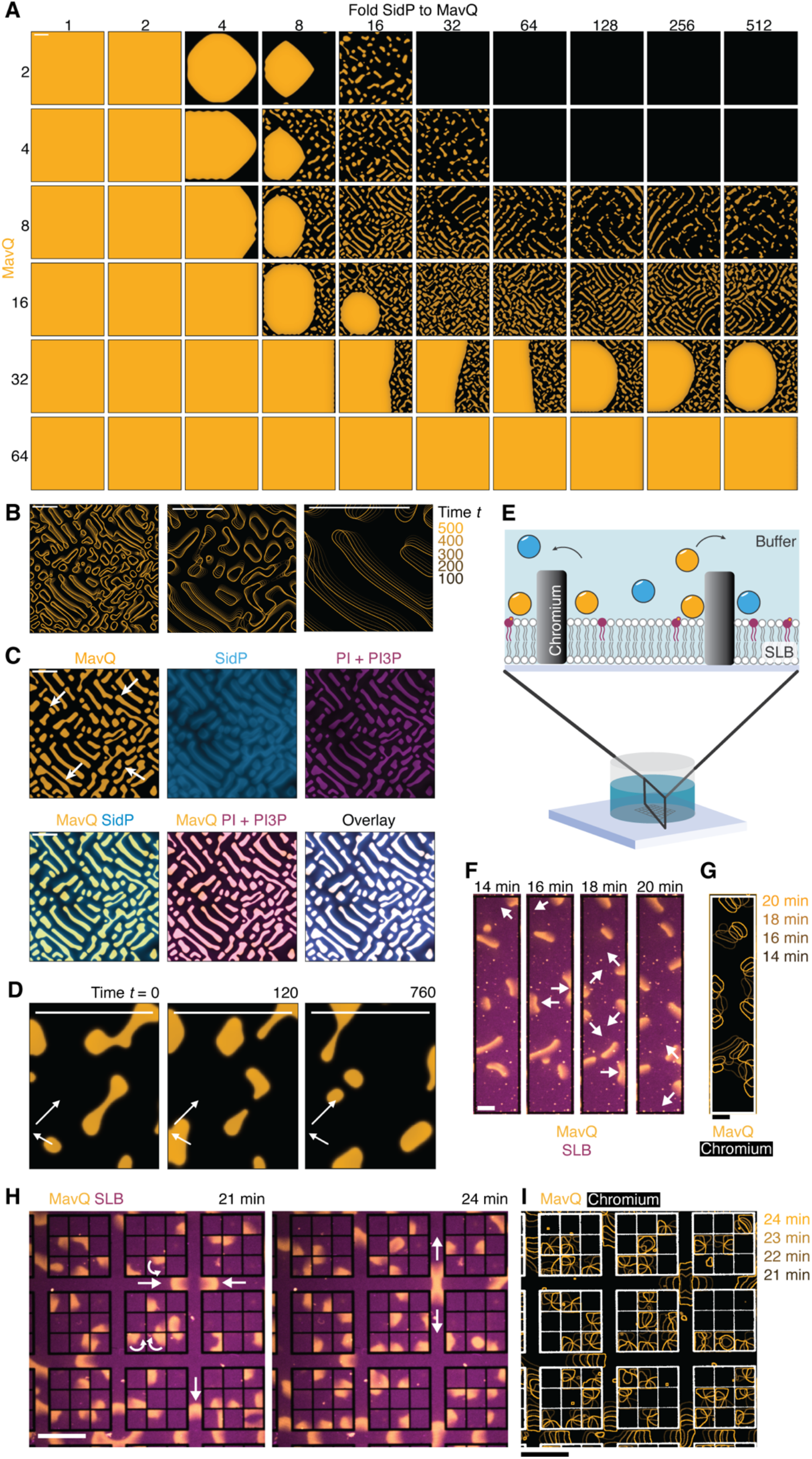
A reaction–diffusion model captures pattern dynamics and predicts geometry-dependent wave propagation. (**A**) Representative snapshots showing the membrane-bound MavQ concentration profile in the reaction–diffusion model at varying bulk MavQ concentrations (*M*) and SidP-to-MavQ ratios (*P*/*M*). The average PI + PI3P lipid area fraction ⟨*φ*⟩ = 0.15. See also movie S20. (**B** and **C**) Pattern contours (B) and concentration profiles (C) for a representative dynamic state (*M* = 8 and *P*/*M* = 16). Each panel in (B), except the first, provides a close-up view of the preceding one. Arrows in (C) indicate the direction of wave propagation. **(D)** Time-series snapshots of membrane-bound MavQ concentration illustrating wavefront reflection at a membrane boundary. Arrows indicate the incident and reflected directions. **(E)** Schematic representation of the in vitro reconstitution assay on geometrically patterned membranes. **(F)** Time-lapse confocal images of MavQ pattern formation on a rectangular membrane patch showing wave reflection at the boundaries. Arrows indicate the direction of wave propagation. See also movie S22. **(G)** Color-coded overlay of pattern contours from the time-series in (F) illustrating wave reflection at the boundaries. **(H)** Time-lapse confocal images of MavQ self-organization on small square corrals and narrow lanes. No coherent patterns emerge across adjacent corrals. Instead, MavQ waves rotated within individual corrals and propagated only along the connecting lanes. Arrows indicate the direction of wave propagation. See also movie S23. **(I)** Color-coded overlay of pattern contours corresponding to (H), showing the absence of coherent patterns across adjacent corrals. Initial lipid composition: 95% DOPC, 5% 18:1 PI, and an extra 0.01% ATTO 647N PE (F to I) Proteins: 20 nM MavQ and 1 µM SidP (F and G); 50 nM MavQ and 2 µM SidP (H and I). For all experiments, MavQ was doped with 25% mEGFP-MavQ, and ATP was added immediately before the start of image acquisition. Scale bars: 100 dimensionless units (A to D); 20 µm (F and G) and 50 µm (H and I).

We next asked whether conservation of the lipid substrates influences pattern dynamics on physically separated membrane patches. Previous work showed that MinDE protein waves, which use the membrane as a passive platform, can couple across thin membrane boundaries through protein diffusion in the shared three-dimensional bulk (*23*). In contrast, MavQ and SidP act on PI and PI3P, which are confined to the membrane. We therefore predicted that the patterns would be restricted to physically continuous membrane surfaces. Consistent with this prediction, simulations demonstrated that MavQ-dense wavefronts were reflected at physical barriers (**Fig. 4D**), whereas MinDE waves propagate across such barriers (*23*). Thus, the model predicts that conservation of the lipid substrates couples MavQ–SidP pattern propagation to membrane geometry.

To test this prediction, we performed in vitro reconstitution assays on geometrically patterned membranes, in which SLBs were partitioned into isolated patches by thin chromium barriers while sharing a common bulk solution (**Fig. 4E**) (*24*). In large rectangular membrane patches, MavQ waves were reflected at the boundaries (**Fig. 4, F and G, and movie S22**), consistent with the model. In smaller square corrals separated by narrow lanes, no coherent patterns emerged across adjacent corrals; instead, MavQ waves rotated within individual corrals and propagated only along the connecting lanes (**Fig. 4, H and I, and movie S23**). Together, these results demonstrate that, unlike MinDE protein waves (*23*), MavQ patterns are confined to physically continuous membrane surfaces, verifying the model prediction that lipid conservation couples pattern propagation to membrane geometry and distinguishes enzyme-driven, substrate-conserving self-organization from systems without such conservation relations on the membrane.

## Discussion

Intracellular patterns arising from the interactions of diffusing proteins are a hallmark of biological self-organization. Our reconstitution of the MavQ–SidP system establishes a minimal in vitro system that generates dynamic, self-organized patterns, via a mechanism distinct from the quintessential MinDE system from *E. coli* (*5*). Unlike MinD and MinE, an ATPase (*25*) and an ATPase-activating protein (*26*), respectively, whose activity cycle drives reversible binding to a charged membrane (*27*), MavQ and SidP form a kinase–phosphatase pair that chemically interconverts PI and PI3P. Thus, dynamic reaction–diffusion-driven patterns can arise not only from reversible membrane binding of proteins but also from enzymatic modification of membrane lipids. The MavQ–SidP system simultaneously patterns both the enzymes and the membrane lipids they modify, integrating lipid chemistry into the pattern-forming mechanism and expanding the repertoire of molecular strategies for intracellular self-organization. Because these lipid substrates are confined to the membrane, wave propagation becomes intrinsically coupled to membrane geometry, while MavQ, SidP, and PI3P remain enriched within the propagating wave itself.

The PI 4-phosphate 5-kinase (PIP5K) and various 5-phosphatases similarly interconvert phosphoinositide species but form bistable spatial patterns in vitro, with at least one enzyme exhibiting cooperative binding to its own reaction product (*13, 28*). As with the MavQ–SidP system, geometric confinement shapes the resulting patterns, and separate confined regions do not correlate with one another. However, PIP5K–5-phosphatase patterns are stationary, arising from intrinsic bistability and stochastic symmetry breaking rather than a propagating front, suggesting that additional interactions, possibly the direct protein–protein interaction between the kinase and phosphatase, as seen in the MavQ–SidP system (*14, 19*), are needed to generate traveling waves. Together, these findings suggest that lipid kinase–phosphatase pairs can pattern membranes through distinct physical mechanisms that give rise to diverse patterning behaviors.

In summary, this work establishes the MavQ–SidP system as a minimal, tractable platform for dissecting how enzymatic modification of membrane lipids generates self-organized patterns, complementing protein-based systems such as MinDE. Extending the system to more complex membrane compositions and additional interacting proteins may reveal how these physical principles operate within phosphoinositide signaling networks. More broadly, cooperative membrane binding coupled to enzymatic feedback inhibition may represent a general design principle for lipid kinase–phosphatase systems acting on a shared membrane substrate, providing a blueprint for engineering synthetic reaction–diffusion circuits (*29–31*) that directly organize membranes as well as proteins.

## Acknowledgments

We thank members of the Tagliabracci, Wingreen, and Ramm laboratories, as well as M. K. Rosen and W. Y. C. Huang, for discussions, and H. Eto and P. Schwille for assistance in the preparation of chromium-patterned coverslips.

## Author contributions

T.-S.H., B.R., and V.S.T. conceived and designed the experiments; T.-S.H. and J.P.P. performed molecular cloning and protein purification; T.-S.H. performed biochemical and cellular experiments; T.-S.H. and B.R. performed imaging experiments; T.-S.H. and B.R. performed image and data analyses with assistance from Y.W. and J.H.; Q.Y. performed theoretical modeling and simulations with input from J.L.Y., Y.M., and N.S.W.; and T.-S.H., B.R., Q.Y., and V.S.T. wrote the manuscript with input from all authors.

## Data and materials availability

All data are available in the main text or the supplementary materials. All materials and code used in this study will be made available upon request.

## Supplementary Materials

### Materials and Methods

#### Chemicals and reagents

Acetic acid (A38212), agar (DF0140), Alexa Fluor 568 NHS ester (A20103), Coomassie brilliant blue G-250 dye (20279), DMEM/High glucose with L-glutamine (11-965-118), glycine (BP381), HCl (A144), HEPES (15630080), HisPur Ni-NTA resin (88221), LB broth (BP97235), NaCl (BP358), sodium pyruvate (11-360-070), SuperSignal West Pico PLUS chemiluminescent substrate (34579), and Tris base (BP152) were obtained from Fisher Scientific (Waltham, MA).

30% Acrylamide/bis solution (1610158), APS (1610700), extra thick blot filter paper (1703966), nitrocellulose membrane (1620112), TEMED (1610801), and Trans-Blot Turbo 5x transfer buffer (10026938) were obtained from Bio-Rad (Hercules, CA).

Agarose (A20090) and glycerol (G22025) were obtained from Research Products International (Mount Prospect, IL).

Ampicillin sodium (A-301), kanamycin monosulfate (K-120), and PMSF (P-470) were obtained from Gold Biotechnology (St. Louis, MO).

AMPPCP (NU-422) was obtained from Jena Bioscience (Jena, Germany).

AMPPNP (A2647), ATP (A9187), ATTO 647N labeled 1,2-dioleoyl-sn-glycero-3-phosphoethanolamine (ATTO 647N PE; 42247), bromophenol blue sodium (B8026), chloramphenicol (C0378), CaCl_2_ (C4901), chloroform (1.02447), EDTA (E5134), ethanol (1.00983), FBS (F2442), Hellmanex III (Z805939), H_2_O_2_ (1.07298), H_2_SO_4_ (339741), imidazole (I2399), IPTG (I5502), KCl (P9541), 2-mercaptoethanol (BME; M3148), methanol (1.06002), MgCl_2_ (M2670), penicillin-streptomycin (P0781), 2-propanol (190764), SDS (L5750), TCEP (C4706), Trolox (238813), and TWEEN 20 (P1379) were obtained from MilliporeSigma (St. Louis, MO).

AZ 351 B (1000351) and AZ ECI 3027 (1A003027) were obtained from MicroChemicals (Ulm, Germany).

1,2-Dioleoyl-sn-glycero-3-phosphocholine (DOPC; 850375), 1,2-dioleoyl-sn-glycero-3-phospho-(1’-myo-inositol) (18:1 PI; 850149), 1,2-dioleoyl-sn-glycero-3-phospho-(1’-myo-inositol-3’-phosphate) (18:1 PI3P; 850150), 1,2-dioleoyl-sn-glycero-3-phospho-L-serine (DOPS; 840035), TopFluor TMR PC (810180), and TopFluor TMR PI (810188) were obtained from Avanti Polar Lipids (Alabaster, AL). PI and PI3P were protonated with HCl before use. Briefly, the lipid powder was dissolved in chloroform to ∼2.5 mM. The solvent was then evaporated under a strong stream of argon, and the resulting lipid film was desiccated under vacuum for 1 h. The dried lipids were redissolved in a 200:100:1 (v/v/v) mixture of chloroform, methanol, and 1 N HCl and incubated for 15 min. The lipids were dried again under argon and desiccated for 1 h. The dried lipids were redissolved in a 3:1 (v/v) chloroform:methanol mixture, dried under argon, redissolved in chloroform, and stored at −20°C.

Ethyl alcohol 200 proof (100129) was obtained from Pharmco (Toronto, Canada).

Gibson Assembly Master Mix (E2611), T4 DNA ligase (M0202), Q5 high-fidelity 2X master mix (M0492), and all restriction enzymes used for cloning were obtained from New England Biolabs (Ipswich, MA).

HCl (97064) and NaOH (M137) were obtained from Avantor (Radnor, PA).

Methoxy PEG silane (M-SLN-5000) was obtained from JenKem Technology (Plano, TX).

Non-functionalized silica microspheres (5-μm silica beads; SS05003) were obtained from Bangs Laboratories (Fishers, IN).

PfuTurbo DNA polymerase (600252) was obtained from Agilent Technologies (Santa Clara, CA).

TransIT-X2 transfection reagent was obtained from Mirus Bio (Madison, WI).

#### Antibodies

Horseradish peroxidase (HRP)-conjugated donkey anti-rabbit antibody (NA934) was obtained from Cytiva (Marlborough, MA). Rabbit anti-MavQ and anti-SidP antibodies (Cocalico Biologicals, Reamstown, PA) were raised against purified MavQ^1–580^ and 6xHis-tagged SidP.

#### Plasmids

*Legionella pneumophila* MavQ and SidP coding sequences (CDS) were amplified by PCR from the genomic DNA of *L. pneumophila* Philadelphia 1 strain Lp02. Amino acid mutations were introduced via QuikChange site-directed mutagenesis. Briefly, primers were designed using the Agilent QuikChange Primer Design Program and used in PCR to generate the desired mutation using PfuTurbo DNA polymerase. Reaction products were digested with Dpn1 and mutations were confirmed by Sanger sequencing.

The wild-type MavQ CDS used for protein expression and purification was the *L. pneumophila* MavQ^1–853^, which lacked the putative C-terminal translocation signal (the C-terminal 19 residues). All MavQ constructs (including wild-type, truncations, mutants, and fluorescent protein-tagged recombinants) were cloned into ppSumo, a modified pet28a bacterial expression vector containing an N-terminal 6xHis tag followed by the yeast Sumo (smt3) CDS. All SidP constructs used for protein expression and purification were cloned into pProEX2, a bacterial expression vector containing an N-terminal 6xHis tag followed by a TEV protease cleavage site. mTagBFP2-2xFYVE_Hrs_ was amplified by PCR from a mammalian expression plasmid (*14*) and cloned into pProEX2 and ppSumo.

The mammalian expression plasmids encoding GFP-MavQ, mCherry-SidP, mTagBFP2-ER, and mTagBFP2-2xFYVE_Hrs_ were previously described (*14*).

All newly generated constructs were verified by Sanger sequencing or whole plasmid sequencing.

#### Bacterial strains, cell lines and culture media

*Escherichia coli* strains were grown in LB broth or on LB agar plates supplemented with 100 μg/mL ampicillin, 34 μg/mL chloramphenicol, or 50 μg/mL kanamycin when appropriate.

HeLa cells were cultured in DMEM/High glucose with L-glutamine supplemented with 1 mM sodium pyruvate, 10% FBS, and 1% penicillin-streptomycin. Cells were maintained at 37°C in a humidified incubator with 5% CO_2_.

#### Mammalian cell transfection and imaging

HeLa cells were seeded on 8-well chambered coverglass (C8-1.5H-N, CellVis, Mountain View, CA) at a density of approximately 2×10^4^ cells/well 1 day before transfection. Cells were transfected with plasmid DNA using TransIT-X2 according to the manufacturer’s instructions. The following amounts of plasmid DNA were used per well: 40 ng of GFP-MavQ, 40-60 ng of mCherry-SidP, 25 ng of mTagBFP2-ER, or 25 ng of mTagBFP2-2xFYVE_Hrs_. Approximately 16 h post-transfection, cells were washed once with extracellular buffer [20 mM HEPES (pH 7.4), 125 mM NaCl, 5 mM KCl, 1.5 mM MgCl_2_, 1.5 mM CaCl_2_, 10 mM glucose] and then equilibrated in extracellular buffer at room temperature for 10 min. Imaging was performed at room temperature using spinning disk confocal microscopy, as detailed in the “Fluorescence microscopy” section below.

#### Protein expression and purification

Plasmids were transformed into Rosetta (DE3) *E. coli* cells. The cells were grown in LB broth at 37°C with shaking until the culture reached an OD_600_ of 0.6-0.8. Protein expression was induced by the addition of 0.4 mM IPTG, and cultures were incubated overnight at 16°C. Cells were harvested by centrifugation and lysed by sonication in lysis buffer [50 mM Tris-HCl (pH 8), 300 mM NaCl, 1 mM PMSF, 14.2 mM BME]. The resulting lysate was clarified by centrifugation at 35,000 × g for 30 min at 4°C. The cleared supernatant was incubated with washed Ni-NTA resin for 1 h at 4°C with gentle rotation. The resin was then loaded onto a gravity-flow column and washed with 80 column volumes of wash buffer [50 mM Tris-HCl (pH 8), 300 mM NaCl, 25 mM imidazole, 5 mM BME]. The protein was eluted with elution buffer [50 mM Tris-HCl (pH 8), 300 mM NaCl, 300 mM imidazole, 5 mM BME]. The 6xHis-SUMO tag was cleaved from the eluted protein by incubation with 6xHis-tagged ULP1 SUMO protease overnight at 4°C. The protein was then further purified by size exclusion chromatography using either a HiLoad Superdex 75 pg or 200 pg column (Cytiva, Marlborough, MA) on an NGC (Bio-Rad, Hercules, CA) or ÄKTA pure chromatography system (Cytiva, Marlborough, MA) in size exclusion buffer [50 mM Tris-HCl (pH 8), 300 mM NaCl, 5 mM TCEP, 10% glycerol].

#### Protein labeling

The amine-reactive dye was dissolved in DMSO to 10 mg/mL. The protein to be labeled was buffer exchanged into labeling buffer [50 mM HEPES (pH 7.5), 300 mM NaCl, 10 mM TCEP, 10% glycerol]. Labeling was carried out by slowly mixing the dye solution into the protein solution at a dye:protein molar ratio of ∼20:1 and then incubated overnight at 4°C. Dye-conjugated proteins were separated using an Econo-Pac 10DG desalting column (Bio-Rad, Hercules, CA).

#### Chromium-patterned coverslips

Chromium-patterned coverslips were fabricated using photolithography and metal evaporation as previously described (*22*). Briefly, coverslips were first rinsed with absolute ethanol and deionized water, followed by drying under a stream of nitrogen. The coverslip surfaces were then activated with oxygen plasma in a Zepto plasma cleaner (Diener Electronic, Ebhausen, Germany) at 40-50% power and 0.3 mbar for 20-60 s. To promote adhesion, the cleaned coverslips were exposed to the vapor of Bis(trimethylsilyl)amine (HDMS) for 2 min. The positive photoresist AZ ECI 3027 was spin-coated onto the HDMS-primed coverslips at 4,000 rpm for 40 s (start/stop acceleration: 2,000 rpm/s). This process yielded a uniform photoresist layer of approximately 3 µm thickness. After pre-baking at 90°C for 90 s, the photoresist was patterned using a µPG101 tabletop micropattern generator (Heidelberg Instruments, Heidelberg, Germany) equipped with a 10 mm write head. The exposure was performed using a 375 nm laser source with a nominal output power of 35 mW before being passed through a 45% attenuation filter. After exposure, the coverslips were post-baked at 110°C for 60 s. The pattern was then developed by immersing the coverslips for 4 min in AZ 351B developer (a NaOH-based solution) diluted 1:4 with deionized water. Finally, the coverslips were rinsed with deionized water and dried under a stream of nitrogen. A chromium layer with a final thickness of ∼30 nm was deposited onto the patterned coverslips by thermal evaporation at a rate of ∼1 Å/s (chamber current: 22-33 mA). For the lift-off procedure, the coverslips were sonicated in acetone for 5 min to remove the remaining photoresist. The coverslips were then rinsed with isopropanol and dried under a stream of nitrogen to yield the final chromium-patterned coverslips.

#### Supported lipid bilayer (SLB) formation

SLBs were formed by fusion of small unilamellar vesicles (SUVs) on cleaned supporting substrates. The methods used were similar to previously published procedures, which are available as a step-by-step video protocol (*32*).

Glass vials (03-391-7B or 03-452-544, Fisher Scientific, Waltham, MA) used for vesicle preparation were cleaned by bath sonication in 3 M NaOH for 45 min, followed by four rinses in deionized water. The vials were then soaked in 5% Hellmanex III at 50°C for 1 h, rinsed 15 times with deionized water, and dried in an oven at 200°C for at least 3 h.

SUVs were prepared by extrusion. A lipid mixture consisting of indicated lipid composition and molar ratios was prepared from chloroform stock solutions in a cleaned glass vial. The solvent was then evaporated under a strong stream of argon, and the resulting lipid film was desiccated under vacuum for 1 h. The lipid film was rehydrated in a rehydration buffer [50 mM Tris-HCl (pH 7.5), 150 or 300 mM KCl], vortexed for 15 min, and extruded 21 times through a 0.05-μm PC membrane using a mini-extruder (Avanti Polar Lipids, Alabaster, AL).

To form SLBs on #1.5H glass coverslips (10812, ibidi, Fitchburg, WI), coverslips were first rinsed with absolute ethanol and deionized water, then dried under a stream of nitrogen. The dried coverslips were etched for 30 min in a freshly prepared piranha solution [H_2_SO_4_:H_2_O_2_ (30%) at a volume ratio of 3:1]. Following etching, the coverslips were thoroughly rinsed with deionized water, dried again under a stream of nitrogen, and affixed to 18-well bottomless sticky slides (81818, ibidi, Fitchburg, WI). 70 μL of the SUV suspension (0.53 mg/mL total lipid) was added to each well and incubated for 30 min at room temperature. Excess vesicles were removed by washing each well 13 times via gentle pipetting in and out 150 μL of rehydration buffer each time.

To form SLBs on chromium-patterned coverslips, coverslips were soaked in 2% Hellmanex III for 30 min at room temperature, rinsed five times with deionized water, and then soaked in a 1:1 (v/v) mixture of isopropanol and deionized water, rinsed 5 times with deionized water, and then dried under a stream of nitrogen. The dried coverslips were etched for 30 min in a freshly prepared piranha solution [H_2_SO_4_:H_2_O_2_ (30%) at a volume ratio of 3:1]. Following etching, the coverslips were thoroughly rinsed with deionized water, dried again under a stream of nitrogen, and affixed to a custom-made round plastic chamber using an optical adhesive (NOA 68, Thorlabs, Newton, NJ). 70 μL of the SUV suspension (0.53 mg/mL total lipid) was added to each chamber and incubated for 30 min at room temperature. Excess vesicles were removed by washing each well 13 times via gentle pipetting in and out 150 μL of rehydration buffer each time.

To prepare the silica bead-supported lipid bilayers (BSLBs), silica beads were first cleaned by etching in 70% nitric acid for at least 2 h. After etching, the beads were collected by centrifugation at 3,428 × g for 5 min. To remove residual acid, the beads were washed with deionized water and re-collected by centrifugation; this wash step was repeated four times. The clean silica beads were then resuspended in rehydration buffer [50 mM Tris-HCl (pH 7.5), 150 or 300 mM KCl] to make a 0.16% (w/v) slurry. After a brief bath sonication, this slurry was mixed with the SUV suspension (1 mM total lipid) at a 5:1 volume ratio, and the mixture was vortexed for 30 min. To remove excess vesicles, the beads were collected by centrifugation at 500 × g for 5 min and washed with rehydration buffer; this wash step was repeated four times.

#### In vitro reconstitution on SLBs

Recombinant proteins were centrifuged at 21,300 × g for 30 min at 4°C, and the soluble supernatant was collected immediately before use.

The reconstitution reaction was assembled on an SLB-coated coverslip in a total volume of 100 µL in reaction buffer [50 mM Tris-HCl (pH 7.5), 150 mM KCl, 5 mM MgCl_2_, 5 mM TCEP, 1 mM Trolox]. Each reaction contained proteins at the concentrations specified and was supplemented with 1 mM ATP where indicated. Image acquisition was initiated either immediately before or immediately after mixing in the final components.

#### Membrane binding assay using BSLBs

#1.5H glass coverslips (10812, ibidi, Fitchburg, WI) were passivated with PEG before use to reduce non-specific protein adsorption in the assay. Coverslips were first rinsed with absolute ethanol and deionized water, then dried under a stream of nitrogen. The dried coverslips were etched for 30 min in a freshly prepared piranha solution [H_2_SO_4_:H_2_O_2_ (30%) at a volume ratio of 3:1]. Following etching, the coverslips were thoroughly rinsed with deionized water, dried again under a stream of nitrogen. The etched coverslips were incubated with a solution of 100 mg/mL methoxy-PEG-silane (MW 5,000) in absolute ethanol at 60°C for at least 5 h. After incubation, the coverslips were thoroughly rinsed with absolute ethanol, followed by deionized water, and dried under a stream of nitrogen. The PEG-passivated coverslips were stored in a vacuum desiccator in the dark until use.

Immediately before an assay, a coverslip was affixed to an 18-well bottomless sticky slide (81818, ibidi, Fitchburg, WI) using an optical adhesive (NOA 68, Thorlabs, Newton, NJ). Recombinant proteins were centrifuged at 21,300 × g for 30 min at 4°C, and the soluble supernatant was collected immediately before use. The binding reaction was assembled in a total volume of 100 µL in reaction buffer [50 mM Tris-HCl (pH 7.5), 150 mM KCl, 5 mM TCEP, 1 mM Trolox]. Each reaction contained proteins at the specified concentrations and approximately 3 µL of a 0.8% (w/v) slurry of BSLBs. Where specified, the reaction was supplemented with 5 mM MgCl_2_ and 1 mM of either ATP, AMPPNP, or AMPPCP. To measure saturation binding, reactions were incubated at room temperature for 4 h to reach equilibrium before being imaged. To monitor binding kinetics, imaging was started immediately after the reaction components were mixed.

#### Liposome co-sedimentation assay

Large unilamellar vesicles (LUVs) were prepared by extrusion. A lipid mixture consisting of indicated lipid composition and molar ratios was prepared from chloroform stock solutions in a cleaned glass vial. The solvent was then evaporated under a strong stream of argon, and the resulting lipid film was desiccated under vacuum for 1 h. The lipid film was rehydrated in a rehydration buffer [50 mM Tris-HCl (pH 7.5), 150 mM KCl], vortexed for 15 min, and then extruded 41 times through a 0.4-μm PC membrane using a mini-extruder (Avanti Polar Lipids, Alabaster, AL).

To remove potential aggregates, recombinant proteins were centrifuged at 21,300 × g for 30 min at 4°C, and the soluble supernatant was collected immediately before use. For the binding reaction, 1 µM of protein was incubated with LUVs (final phosphoinositide concentration of 150 µM) in a total volume of 80 µL of reaction buffer [50 mM Tris-HCl, pH 7.5, 150 mM KCl, 5 mM TCEP, 1 mM Trolox]. The reactions were incubated for 4 h at room temperature and then centrifuged at 20,000 × g for 30 min at 4°C to pellet the liposomes. The supernatant was collected, mixed with 20 µL of 5x SDS-PAGE loading buffer, and boiled. The liposome pellet was resuspended in 100 µL of 1x SDS-PAGE loading buffer and boiled. Equal volumes of the supernatant and pellet fractions were resolved by SDS-PAGE. Protein distribution was visualized by western blotting and quantified by densitometry.

#### Fluorescence microscopy

Spinning disk confocal microscopy and total internal reflection fluorescence (TIRF) microscopy were performed on an Olympus IXplore SpinSR10 system equipped with a cellTIRF module (Olympus, Waltham, MA). Spinning disk confocal images were acquired using a U Plan S-Apo 40x/0.95 objective (Olympus, Waltham, MA) and an Andor iXon Ultra 888 electron multiplier charge-coupled device (EM-CCD) camera (Oxford Instruments, Belfast, UK). TIRF images were acquired using an Apo N 60x/1.49 TIRFM oil objective (Olympus, Waltham, MA) and an Andor iXon Ultra 888 EM-CCD camera.

#### Image, data and statistical analyses

Fluorescence images were displayed using the blue, orange, and purple lookup tables (BOP LUT scheme; https://github.com/cleterrier/ChrisLUTs). Data visualization, statistical analyses, and curve fitting were performed using GraphPad Prism 10 (GraphPad Software, La Jolla, CA), unless otherwise specified.

To visualize wave propagation, wave patterns were segmented and temporally color-coded using a custom ImageJ/Fiji macro. First, raw fluorescence images were smoothed using a median filter (radius: 10 pixels). A pseudo-flat-field correction (BioVoxxel Toolbox; radius: 300 pixels) was subsequently applied to correct for inhomogeneous illumination. The corrected images were thresholded using the Huang method to generate distinct wave outlines. Following a dilation operation, the wave pattern outlines were color-coded using the ZstackDepthColorCode v0.0.2 plugin (https://github.com/UU-cellbiology/ZstackDepthColorCode) and rendered as a final maximum intensity projection.

For spatial line profiles, images were smoothed using the Gaussian blur function (sigma: 5 pixels) of ImageJ/Fiji. To eliminate background from the SidP channel for both visualization and spatial line profiles, a sliding window average calculated over 60 frames was subtracted from the raw time series.

Wave velocities were quantified by tracking individual wavefronts. The trailing edges of the moving waves were manually tracked using the mTrackJ plugin for ImageJ/Fiji, and the average velocity was calculated for each independent track. For each analyzed image stack, 2–3 distinct waves were tracked.

To analyze average fluorescence intensities and the coefficient of variation, bright protein aggregates and local membrane defects were computationally filtered and removed using the Remove Outliers function of ImageJ/Fiji before quantification.

Kymographs were generated to track spatiotemporal dynamics using the Multi Kymograph plugin for ImageJ/Fiji. Space-time plots were extracted along the 20-pixel-wide linear selection indicated in the corresponding figure panel.

To analyze MavQ binding and PI3P production on BSLBs, a custom ImageJ/Fiji macro was used. A binary mask defining the BSLB area was generated by applying the default dark auto-thresholding method to the BSLB channel. To account for background fluorescence, a fixed 50 × 50 pixel squared region of interest (ROI) was specified in a region containing no beads or protein aggregates, and the mean background intensity was recorded for both channels. The mean fluorescence intensities within the masked BSLB areas were then quantified, and their respective background values were subtracted. To quantify relative membrane binding, the background-subtracted MavQ or 2xFYVE_Hrs_ fluorescence intensity was normalized by dividing it by the corresponding background-subtracted BSLB intensity.

To visualize and enhance the contrast of labeled lipids, time-series fluorescence image stacks were processed using custom Python scripts. To isolate local, transient changes in lipid intensity from slow background or illumination drift, a rolling background subtraction routine was implemented. For each frame in a given stack, background was calculated as the 55th percentile intensity of the preceding 10 frames. Both the current frame and its corresponding background were spatially smoothed using a Gaussian filter (sigma: 2 pixels). The smoothed background was then subtracted from the smoothed current frame to yield a localized difference map. Depending on the visualization requirements, the difference maps were processed using two distinct pipelines:

1. To selectively isolate regions of local lipid enrichment, negative intensity values in the difference map were clipped to zero, retaining only positive brightness fluctuations.
2. To capture both local lipid enrichment and depletion zones, the full range of both positive and negative intensity values was preserved.

To ensure consistent brightness and contrast across the entire time series, a global intensity normalization protocol was applied. All difference maps across all frames were compiled into a single collective distribution, and the 1st and 99th percentiles were calculated to establish fixed lower and upper normalization thresholds, respectively. Pixel intensities across all frames were linearly scaled between these global bounds and clipped to an interval of [0, 1]. The normalized values were rendered by mapping the scalar values to a blue-white-red colormap using Matplotlib.

#### Replication

All experiments were performed at least 3 times unless otherwise stated.

#### Data availability

All the data supporting the findings of this study are available within the paper and its supplemental information files. Other relevant data are available from the corresponding authors upon reasonable request.

### Supplementary Text

#### Continuum model

We consider a phenomenological reaction–diffusion model for MavQ–SidP pattern formation on a supported lipid bilayer. The model describes three fields: The membrane-bound MavQ density *m*(*x*, *t*), the membrane-bound SidP density *p*(*x*, *t*), and the combined PI and PI3P lipid fraction *ϕ*(*x*, *t*). The protein fields are nondimensionalized by the local membrane capacity, such that:

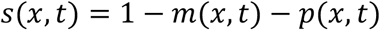

represents the local free membrane capacity available for further binding. The densities of MavQ and SidP evolve according to the following dynamics:

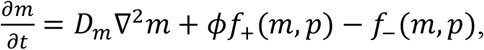

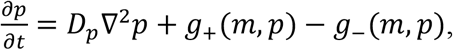

where *D_m_* and *D_p_* are the diffusion coefficients of membrane-bound MavQ and SidP, respectively. The terms *f*_±_(*m*, *p*) represent the binding and unbinding of MavQ, and *g*_±_(*m*, *p*) for SidP. We assume that MavQ can only bind where PI and PI3P lipids are present, such that the binding rate is proportional to the lipid fraction *ϕ*. We also assume the following binding and unbinding kinetics:

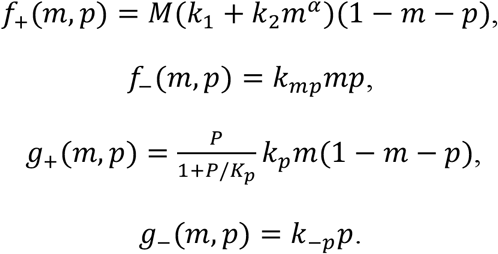

Here, *M* and *P* are the bulk MavQ and SidP concentrations, respectively. *k*_1_ and *k*_2_ are the basal and cooperative binding rates of MavQ, with the exponent *α* controlling the level of cooperativity. *k_mp_* is the rate of MavQ unbinding facilitated by SidP through its PI 3-phosphatase activity. *k_p_* is the rate constant for SidP recruitment facilitated by membrane-bound MavQ. The binding rate saturates at high bulk SidP concentrations with half-maximum at *P* = *K_p_*. The rate constant for SidP dissociation is *k*_−*p*_.

The combined PI and PI3P lipid fraction evolves following a continuity equation:

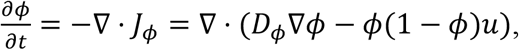

where *D_ϕ_* is the diffusion coefficient of lipids, and *u* is the velocity of lipid transport driven by attractive interactions with MavQ. The local conservation of lipids affects the dynamics of wavefronts, especially near the boundary where lipids cannot diffuse out of the system. Here, we take 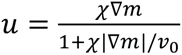, which allows lipids to move up the gradient of membrane-bound MavQ at a velocity that does not exceed *v*_0_. We find that the lipid transport is necessary for pattern formation (**Fig. S7D**). Finally, the mobility factor *ϕ*(1 − *ϕ*) ensures that drift does not drive *ϕ* below 0 or above 1.

This model is intended as a minimal phenomenological description of the patterns. It does not distinguish PI from PI3P explicitly and does not include separate phosphorylation and dephosphorylation reactions. Instead, *ϕ* represents the pooled lipid species that promote MavQ membrane recruitment and that are redistributed by MavQ-dependent transport.

#### Numerical simulations

The simulations were performed in a square domain with no flux boundary conditions. The domain was discretized into an *N* × *N* grid with *N* = 512 and spacing *Δx* = 1. The solver was implemented in JAX. Spatial derivatives were approximated using second-order finite differences, with central stencils in the interior and one-sided stencils at the boundary. Time integration used an explicit predictor–corrector update. For each attempted step, reaction and transport terms were first evaluated at the current state, and then two Picard corrector iterations were performed, in which the reaction term was trapezoidally averaged between the old and predicted states, while diffusion and transport were evaluated at the predicted state. The following adaptive timestepping was used: If the correction norm exceeded 0.1, if any finite field became nonfinite, or if *m* or *p* left the tolerated interval [−0.01, 1.01], the attempted step was rejected and the time step was halved. Accepted steps increased the next time step by a factor of 1.1 up to *Δt_max_* = 0.05.

Unless otherwise stated, all simulations used the parameter values listed in **Table S1** (expressed in nondimensional units). To obtain steady-state phase diagrams, simulations were run for a sufficiently long time (*T* ∼ 2 × 10^6^) from an inhomogeneous initial condition. Additional short simulations were performed to produce the dynamics in the contour plots and supplemental movies.

---

**Table S1.**
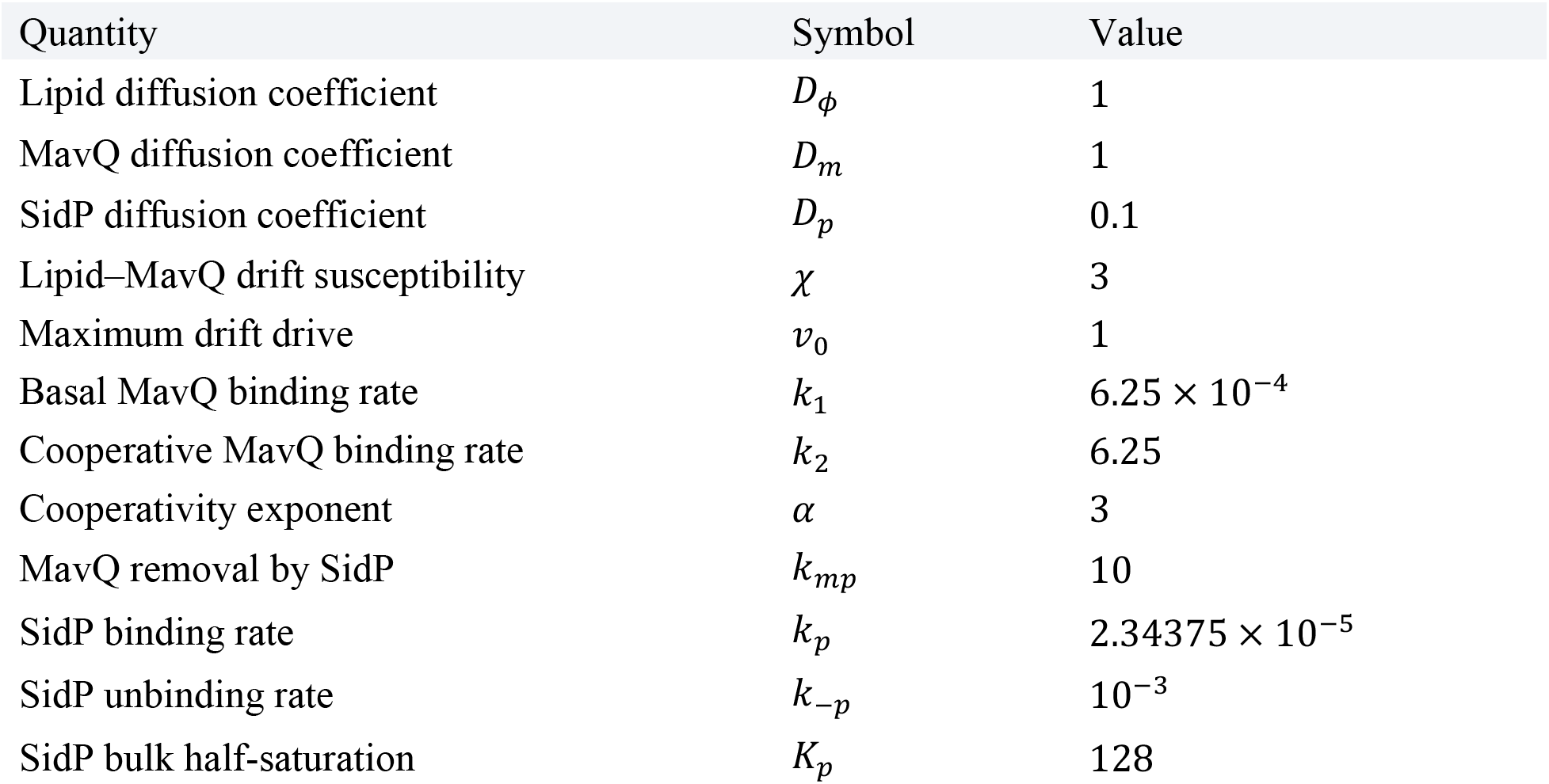
Default model parameters used for the simulations.

**Fig. S1.**
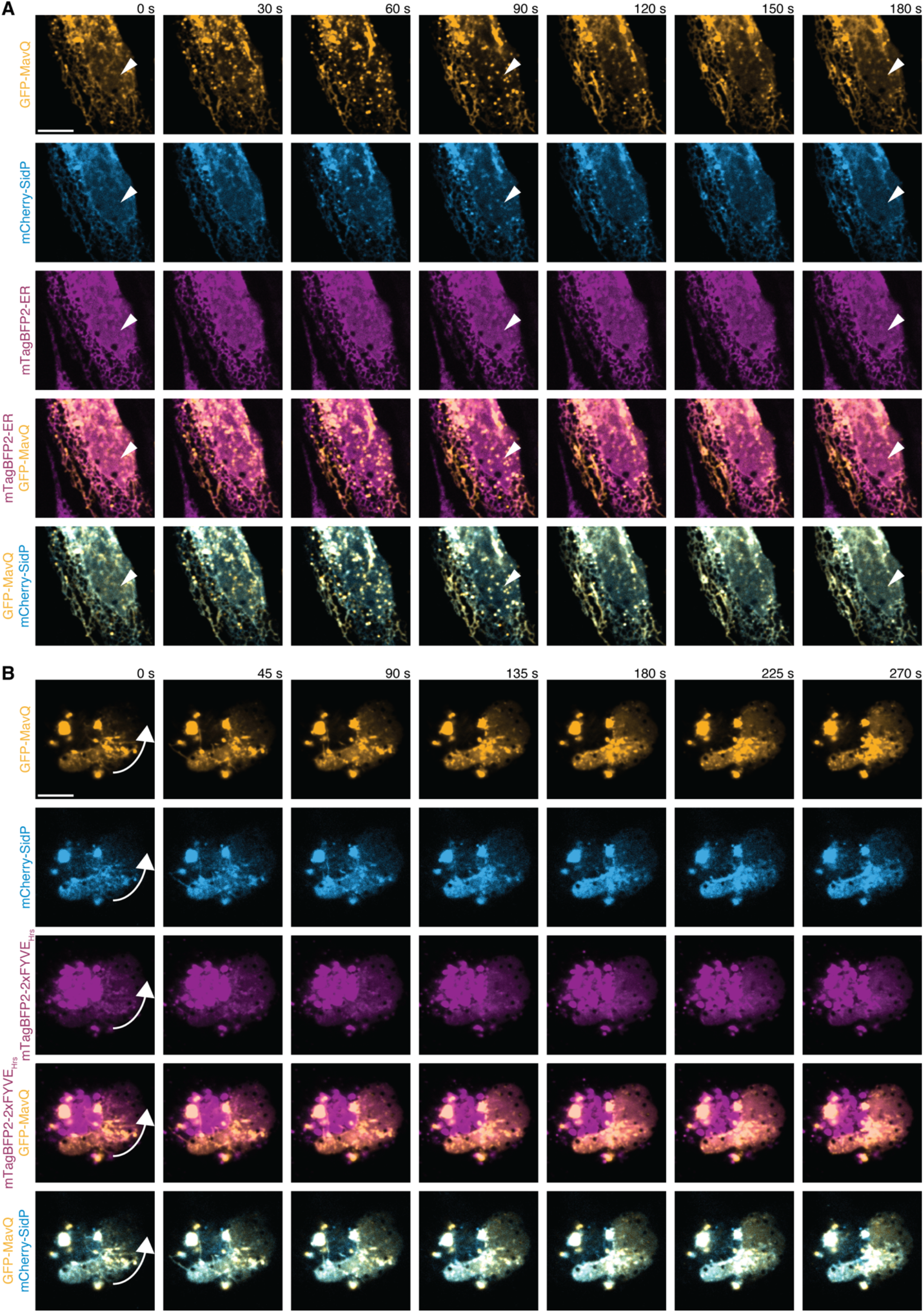
The *Legionella* PI 3-kinase MavQ and the 3-phosphatase SidP generate dynamic protein and phosphoinositide patterns on the ER. **(A)** Confocal images showing the localization of GFP-MavQ and mCherry-SidP relative to the ER in a live HeLa cell. Initially, both proteins localize predominantly to the ER. Between 0 and 90 s, MavQ- and SidP-enriched vesicles emerge. The vesicle-localizing MavQ and SidP then dissipate and redistribute onto and propagate along the ER network. Arrowheads serve as landmarks. See also movie S1 for better visualization. **(B)** Confocal images showing the localization of the PI3P probe mTagBFP2-2xFYVE_Hrs_ relative to GFP-MavQ and mCherry-SidP in a live HeLa cell. A PI3P wave propagates along the perinuclear ER and nuclear envelope, colocalizing with MavQ and SidP. Arrows indicate the direction of wave propagation. See also movie S2 for better visualization. Fluorescence signals were intentionally oversaturated in regions with high MavQ, SidP, or PI3P abundance (vesicular-tubular structures and ER-associated clumps enriched with MavQ and SidP, or PI3P-positive endosomes without MavQ and SidP) to facilitate visualization of the relatively weak signals on the perinuclear ER and nuclear envelope. Scale bars: 10 μm.

**Fig. S2.**
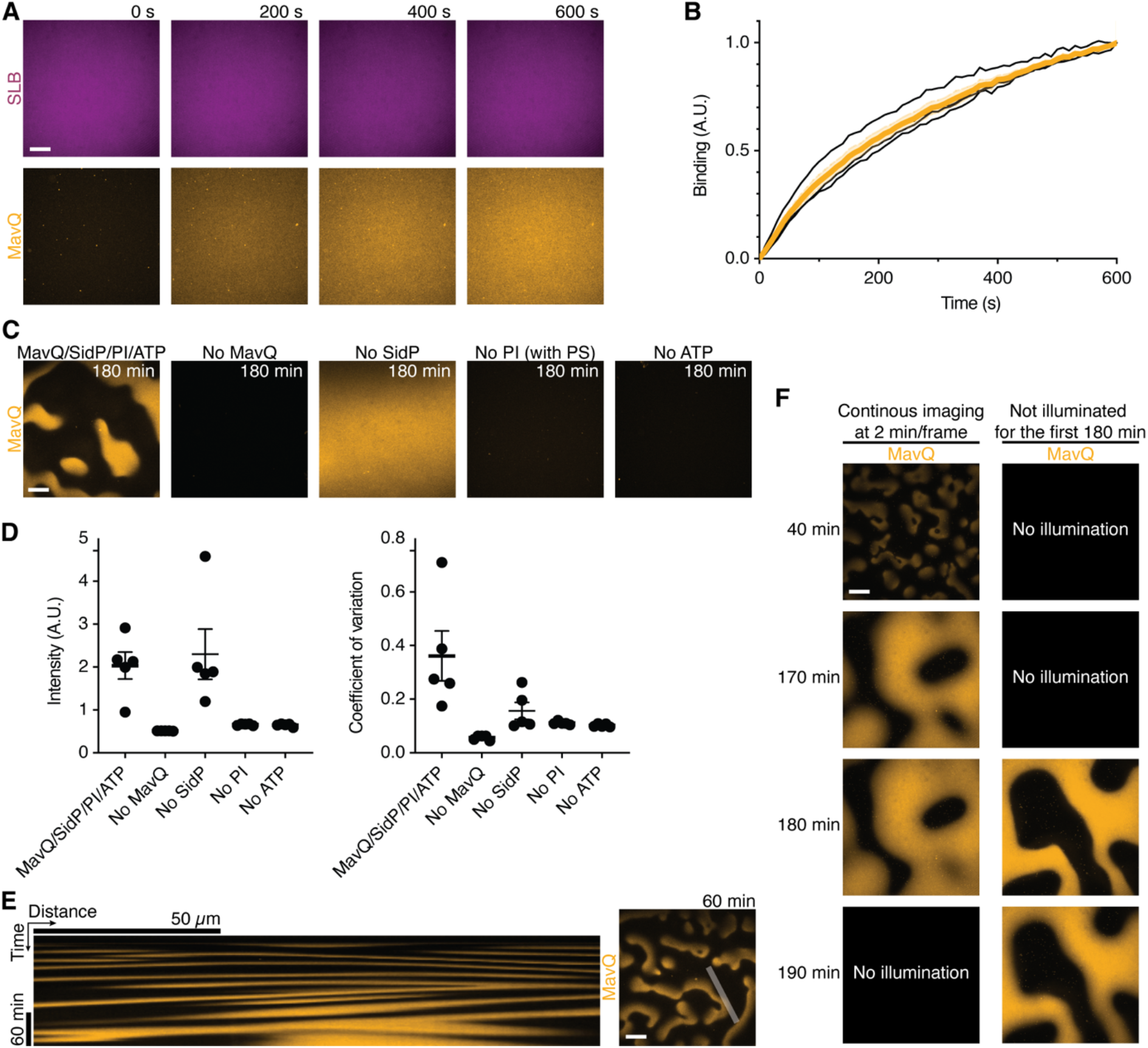
MavQ, SidP, PI and ATP are necessary and sufficient for pattern formation. (**A** and **B**) Time-lapse confocal images (A) and corresponding kinetics (B) of MavQ binding on an SLB. ATP was added immediately before the start of image acquisition. Curves: traces from 4 independent experiments in black; mean and SEM in (B) in orange. **(C)** Confocal images of MavQ intensity on the membrane in reconstitution assays with the indicated components, taken 180 min after ATP addition. The phospholipid phosphatidylserine (PS) was included in the SLB for the “No PI” condition to account for membrane charges. Brightness and contrast are comparable among all images. See also movie S3. **(D)** Quantification of MavQ intensity and its coefficient of variation for the conditions shown in (C), analyzed between 60 and 90 min after ATP addition. Bars: mean and SEM of 5 replicates across 3 independent experiments. **(E)** Kymograph (taken along the gray line marked in a snapshot from Fig. 1B) showing dynamic patterning and wave propagation. See also the first panel of movie S3. **(F)** Control experiment demonstrating that the formation of quasi-stationary patterns is not a phototoxicity artifact. Similar patterns emerged in chambers that were illuminated continuously for 180 min (2 min/frame) or were kept in the dark for 180 min before imaging. See also movie S4. Initial lipid composition: 95% DOPC, 5% 18:1 PI (or substituted with PS as indicated), and an extra 0.01% ATTO 647N PE. Proteins: 10 nM MavQ (A and B), or 10 nM MavQ with 1 µM SidP (C to F). For all panels, MavQ was doped with 25% mEGFP-MavQ. Scale bars: 50 µm.

**Fig. S3.**
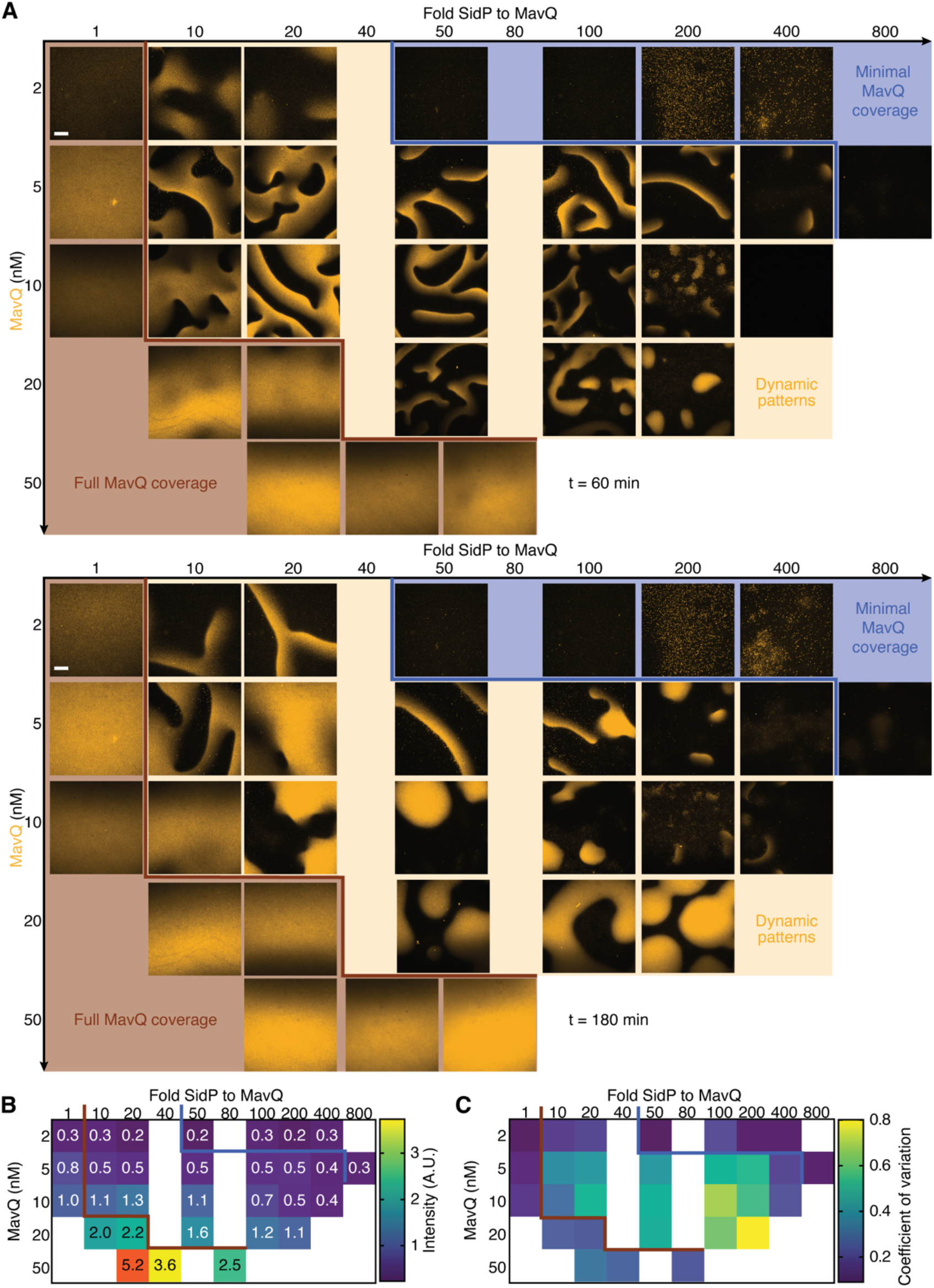
Distinct regimes of spatiotemporal patterns are governed by the MavQ concentration and the SidP-to-MavQ molar ratio. **(A)** Confocal images at two time points illustrating the three patterning regimes as a function of MavQ concentration and the SidP-to-MavQ molar ratio. Brightness and contrast are comparable within each row. See also Fig. 1G and movie S6. (**B** and **C**) Heat maps of the mean MavQ fluorescence intensity (B) and the coefficient of variation of MavQ fluorescence intensity (C), analyzed between 60 and 90 min after ATP addition. Higher coefficient of variation indicates the pattern-forming regime. Data points: mean from 4 independent experiments, with at least 2 repeats per condition. Initial lipid composition: 95% DOPC, 5% 18:1 PI, and an extra 0.01% ATTO 647N PE. Proteins: as indicated in the panels. For all experiments, MavQ was doped with 25% mEGFP-MavQ, and ATP was added immediately before the start of image acquisition. Scale bars: 50 µm.

**Fig. S4.**
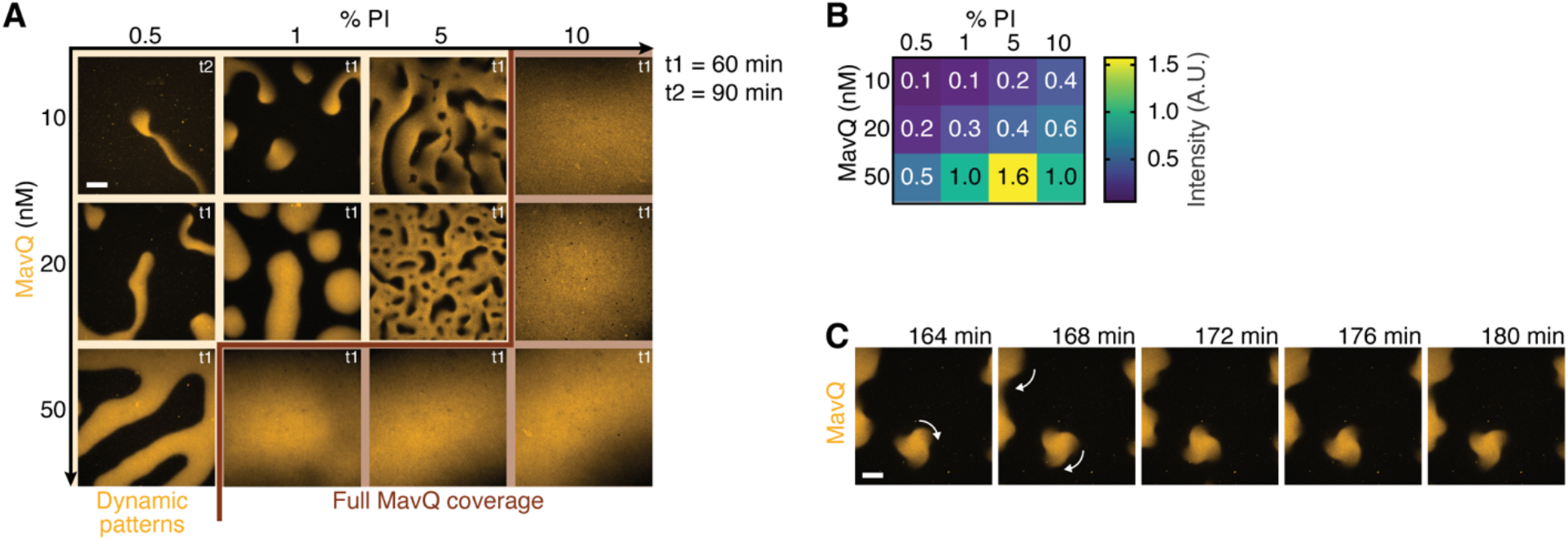
PI concentration in the membrane influences dynamic pattern formation of MavQ. **(A)** Confocal images illustrating how MavQ spatiotemporal patterns are modulated by the concentrations of MavQ and PI. Note that brightness and contrast are not comparable between images. See also movie S7. **(B)** Heat maps of the mean MavQ fluorescence intensity analyzed between 60 and 90 min after ATP addition. Data points: mean of 3 independent experiments. **(C)** Time-lapse confocal images of MavQ spots and stripes with wavy edges. Arrows indicate the direction of rotation. See also movie S7. Initial lipid composition: 90-99.5% DOPC, 0.5-10% 18:1 PI, and an extra 0.01% ATTO 647N PE (A and B); 99% DOPC, 1% 18:1 PI, and an extra 0.01% ATTO 647N PE (C). Proteins: 10-50 nM MavQ (A and B) or 10 nM MavQ (C), each with a 50-fold molar excess of SidP. For all experiments, MavQ was doped with 25% mEGFP-MavQ, and ATP was added immediately before the start of image acquisition. Scale bars: 50 µm.

**Fig. S5.**
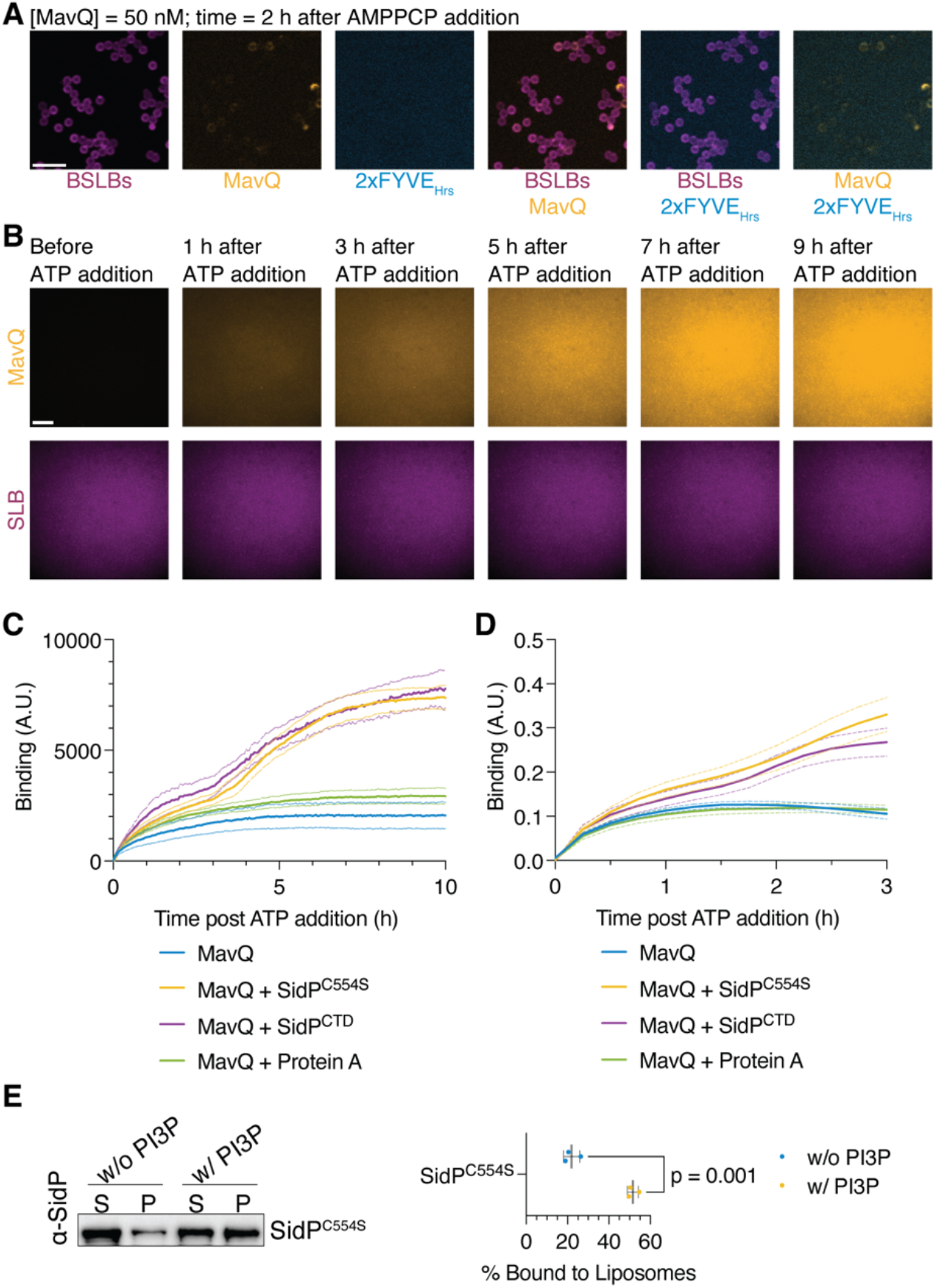
SidP’s PI 3-phosphatase activity provides the negative feedback required for pattern formation. **(A)** Confocal images of MavQ binding and PI3P production on PI-containing BSLBs, taken 2 h after AMPPCP addition. PI3P was visualized with the probe mTagBFP2-2xFYVE_Hrs_. See also movie S9. **(B)** Time-lapse confocal images depicting MavQ membrane binding on a PI-containing SLB in the presence of SidPC^554S^, taken at the indicated time points before and after ATP addition. (**C** and **D**) Time course depicting MavQ membrane binding on PI-containing SLBs (C) or BSLBs (D) in the absence or presence of SidP^C554S^, SidP^CTD^, or Protein A. Curves: mean and SEM of 3 (C) or 4 (D) independent experiments. (**E**) Liposome sedimentation assay depicting the binding of SidP^C554S^ to DOPC or PI3P-containing liposomes. S: supernatant; P: pellet. Note that the inactive SidP was used in the assays to prevent hydrolysis of PI3P. The strip plot quantifies the fraction of protein co-sedimenting with liposomes. Liposome-bound fractions were quantified as the pellet fraction normalized to the total protein. Bars: mean and SD of 3 independent experiments; p-values from two-tailed t-tests. Initial lipid composition: 95% DOPC, 5% 18:1 PI, and an extra 0.01% ATTO 647N PE (A to D); 100% DOPC or 85% DOPC with 15% 18:1 PI3P (E). Proteins: 50 nM MavQ, doped with 25% mEGFP-MavQ, and 50 nM mTagBFP2-2xFYVE_Hrs_ (A); 10 nM MavQ, doped with 25% mEGFP-MavQ, and 1 μM SidP^C554S^ (B); 10 nM MavQ, doped with 25% mEGFP-MavQ, and 1 μM SidP^C554S^, SidP^CTD^, or Protein A. (C and D); 1 μM SidP^C554S^ (E). Scale bars: 20 µm (A); 50 µm (B).

**Fig. S6.**
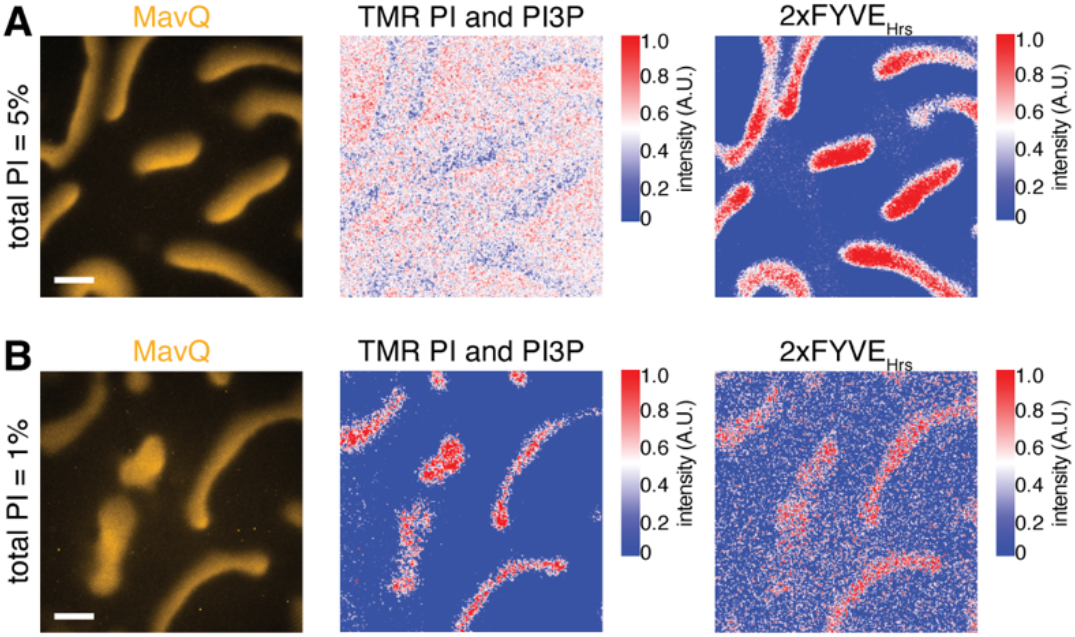
Self-organization of MavQ and SidP drives spatiotemporal patterning of lipid substrates in membranes. (**A**) Confocal images and heat maps showing MavQ patterns and phospholipid distribution in an SLB containing 5% total PI. The combined PI and PI3P pool was tracked using TopFluor TMR PI, while PI3P was specifically visualized using mTagBFP2-2xFYVE_Hrs_. See also movie S13. (**B**) Confocal images and heat maps showing MavQ patterns and the underlying phospholipid distribution in an SLB containing 1% total PI. The combined PI and PI3P pool was tracked using TopFluor TMR PI, while PI3P was specifically visualized using mTagBFP2-2xFYVE_Hrs_. See also movie S17. Initial lipid composition: 95% DOPC, 4.9% 18:1 PI, and 0.1% TopFluor TMR PI (A); 99% DOPC, 0.9% 18:1 PI, and 0.1% TopFluor TMR PI (B). Proteins: 20 nM MavQ, doped with 25% mEGFP-MavQ, 2 µM SidP, and 20 nM mTagBFP2-2xFYVE_Hrs_. ATP was added immediately before the start of image acquisition. Scale bars: 50 µm.

**Fig. S7.**
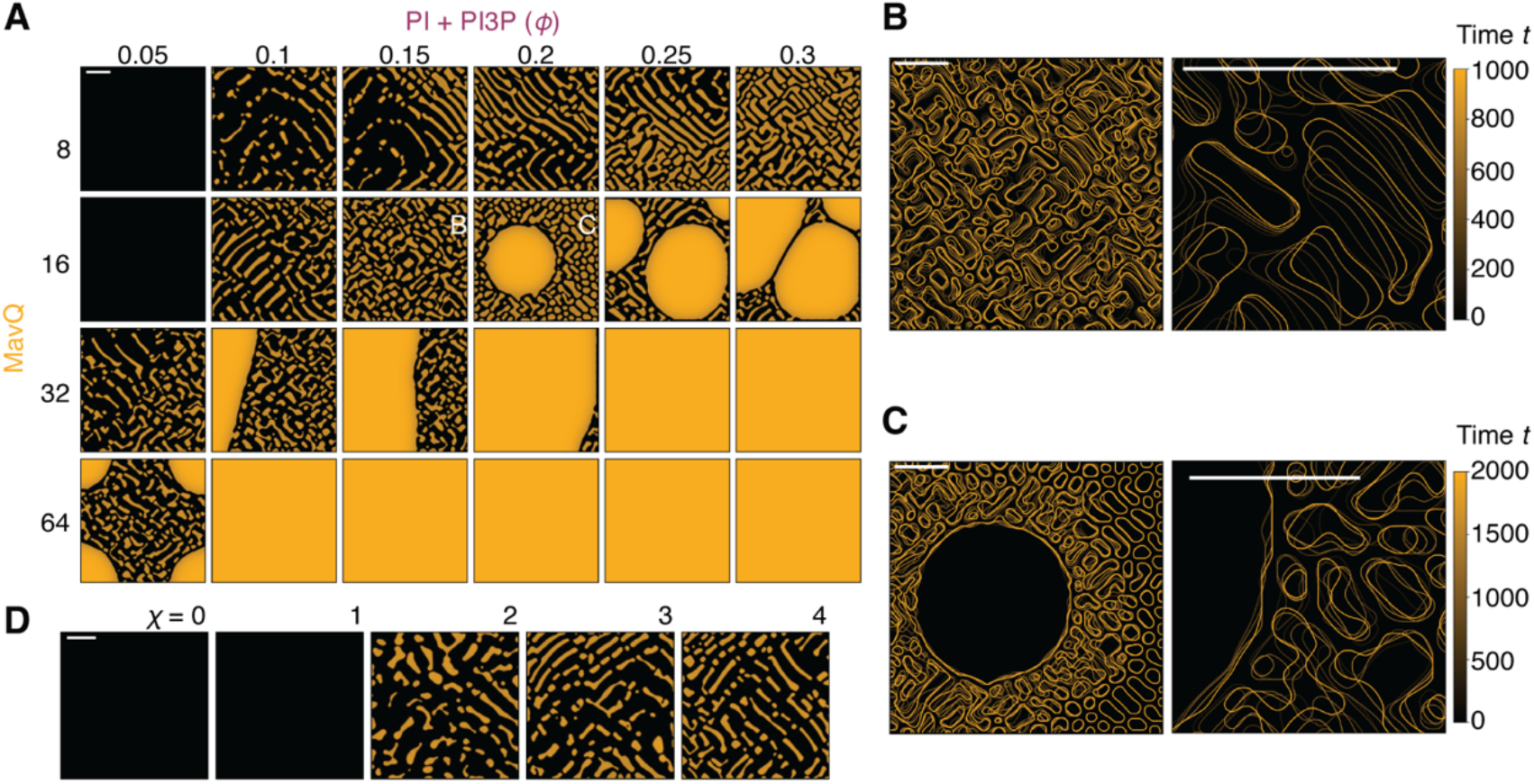
Pattern dynamics depend on MavQ concentration, lipid abundance and lipid transport. (**A**) Representative snapshots of membrane-bound MavQ concentration profile in the reaction– diffusion model at varying bulk MavQ concentrations (*M*) and average PI + PI3P lipid area fractions (⟨*ϕ*⟩). The bulk SidP-to-MavQ ratio was fixed at *P*/*M* = 32. See also movie S21. (**B** and **C**) Pattern contours at successive time points for the two conditions in (A): *M* = 16; ⟨*ϕ*⟩ = 0.15 for (B), and ⟨*ϕ*⟩ = 0.20 for (C). (**D**) Representative snapshots illustrating the dependence of pattern dynamics on MavQ-directed lipid transport at varying values of the Lipid–MavQ drift susceptibility (*χ*). *M* = 32, *P*/*M* = 8, ⟨*ϕ*⟩ = 0.15. Scale bars: 100 dimensionless units.

**Movie S1.**

Time-lapse confocal imaging of GFP-MavQ and mCherry-SidP dynamics relative to the ER in a live HeLa cell. Initially, both proteins distribute mostly on the ER. MavQ- and SidP-enriched vesicles emerge between 0 and 90 s. The vesicle-localizing MavQ and SidP then dissipate and redistribute onto and propagate along the ER network. Scale bar: 10 µm.

**Movie S2.**

Time-lapse confocal imaging showing the dynamics of the PI3P probe mTagBFP2-2xFYVE_Hrs_ relative to GFP-MavQ and mCherry-SidP in a live HeLa cell. A PI3P wave propagates on the perinuclear ER and nuclear envelope, tracking MavQ and SidP. Note that fluorescence signals for MavQ, SidP, and the PI3P probe are oversaturated at highly concentrated regions (vesicular-tubular structures and ER-associated clumps enriched with MavQ and SidP, or PI3P-positive endosomes without MavQ and SidP) to enable visualization of the relatively weak signals on the perinuclear ER and nuclear envelope. Scale bar: 10 µm.

**Movie S3.**

Time-lapse confocal imaging of MavQ intensity on the membrane in reconstitution assays with the indicated components. ATP was added before image acquisition. Initial lipid composition: 95% DOPC, 5% 18:1 PI, and an extra 0.01% ATTO 647N PE. Proteins: 10 nM MavQ, doped with 25% mEGFP-MavQ, and 1 µM SidP. Scale bars: 50 µm.

**Movie S4.**

Time-lapse confocal imaging showing that similar MavQ patterns emerged in chambers that were illuminated continuously for 180 min (2 min/frame) or were kept in the dark for 180 min before imaging. ATP was added before the start of the experiment. Initial lipid composition: 95% DOPC, 5% 18:1 PI, and an extra 0.01% ATTO 647N PE. Proteins: 10 nM MavQ, doped with 25% mEGFP-MavQ, and 1 µM SidP. Scale bars: 50 µm.

**Movie S5.**

Time-lapse total internal reflection fluorescence imaging shows SidP traveling with MavQ within dynamic membrane patterns. For the SidP channel, the static background, estimated by the average intensity across all frames, has been subtracted for clarity. ATP was added before image acquisition. Initial lipid composition: 95% DOPC, 5% 18:1 PI, and an extra 0.01% ATTO 647N PE. Proteins: 10 nM MavQ, doped with 25% mEGFP-MavQ, and 2 µM SidP, doped with 25% Alexa Fluor 568-labeled SidP. Scale bar: 50 µm.

**Movie S6.**

Time-lapse confocal imaging illustrating the distinct regimes of MavQ patterns that are governed by the MavQ and SidP concentrations. ATP was added before image acquisition. Initial lipid composition: 95% DOPC, 5% 18:1 PI, and an extra 0.01% ATTO 647N PE. Proteins: as indicated in the panels. For all panels, MavQ was doped with 25% mEGFP-MavQ. Scale bars: 50 µm.

**Movie S7.**

Time-lapse confocal images illustrating how MavQ spatiotemporal patterns are modulated by the concentrations of MavQ and PI. Note that brightness and contrast are not comparable between images. ATP was added before image acquisition. Initial lipid composition: 90-99.5% DOPC, 0.5-10% 18:1 PI, and an extra 0.01% ATTO 647N PE. Proteins: 10-50 nM MavQ with a 50-fold molar excess of SidP. For all experiments, MavQ was doped with 25% mEGFP-MavQ. Scale bars: 50 µm.

**Movie S8.**

Time-lapse confocal imaging of PI-containing BSLBs incubated with MavQ and a PI3P probe in the presence of ATP, which was added before image acquisition. BSLBs are shown in purple, MavQ in orange, and PI3P (visualized by mTagBFP2-2xFYVE_Hrs_) in blue. Bar: 50 µm. Area shown in Fig. 2A is boxed. The initial lipid composition: 95% DOPC, 5% 18:1 PI, and 0.01% ATTO 647N PE. Proteins: 50 nM MavQ, doped with 25% mEGFP-MavQ, and 50 nM mTagBFP2-2xFYVE_Hrs_.

**Movie S9.**

Time-lapse confocal imaging of PI-containing BSLBs incubated with MavQ and a PI3P probe in the presence of AMPPCP, which was added before image acquisition. BSLBs are shown in purple, MavQ in orange, and PI3P (visualized by mTagBFP2-2xFYVE_Hrs_) in blue. Note that non-specific protein binding to the silica bead is present where the membrane is defected. Bar: 50 µm. Area shown in Fig. S5A is boxed. The initial lipid composition: 95% DOPC, 5% 18:1 PI, and 0.01% ATTO 647N PE. Proteins: 50 nM MavQ, doped with 25% mEGFP-MavQ, and 50 nM mTagBFP2-2xFYVE_Hrs_.

**Movie S10.**

Time-lapse confocal imaging of PI-containing BSLBs incubated with MavQ and a PI3P probe in the presence of AMPPNP, which was added before image acquisition. BSLBs are shown in purple, MavQ in orange, and PI3P (visualized by mTagBFP2-2xFYVE_Hrs_) in blue. Note that MavQ binding and PI3P production occur in an all-or-none fashion. Bar: 50 µm. Area shown in Fig. 2D is boxed. The initial lipid composition: 95% DOPC, 5% 18:1 PI, and 0.01% ATTO 647N PE. Proteins: 50 nM MavQ, doped with 25% mEGFP-MavQ, and 50 nM mTagBFP2-2xFYVE_Hrs_.

**Movie S11.**

Time-lapse confocal imaging showing colocalization of MavQ and PI3P within patterns. PI3P was visualized with the probe mTagBFP2-2xFYVE_Hrs_. ATP was added before image acquisition. Initial lipid composition: 95% DOPC, 5% 18:1 PI, and an extra 0.01% ATTO 647N PE. Proteins: 20 nM MavQ, doped with 25% mEGFP-MavQ, 2 µM SidP, and 20 nM mTagBFP2-2xFYVE_Hrs_. Scale bar: 50 µm.

**Movie S12.**

Time-lapse confocal imaging of MavQ patterns and the underlying PI distribution in an SLB containing 5% total PI. The distribution of PI was tracked using TopFluor TMR PI. ATP was added before image acquisition. Initial lipid composition: 95% DOPC, 4.9% 18:1 PI, and 0.1% TopFluor TMR PI. Proteins: 20 nM MavQ, doped with 25% mEGFP-MavQ, and 2 µM SidP. Scale bar: 50 µm.

**Movie S13.**

Time-lapse confocal imaging of MavQ patterns and the underlying lipid substrate distribution in an SLB containing 5% total PI. The distribution of the combined PI and PI3P pool was tracked using TopFluor TMR PI, while PI3P was specifically visualized using mTagBFP2-2xFYVE_Hrs_. ATP was added before image acquisition. Initial lipid composition: 95% DOPC, 4.9% 18:1 PI, and 0.1% TopFluor TMR PI. Proteins: 20 nM MavQ, doped with 25% mEGFP-MavQ, 2 µM SidP, and 20 nM mTagBFP2-2xFYVE_Hrs_. Scale bar: 50 µm.

**Movie S14.**

Time-lapse confocal imaging of MavQ patterns and the underlying PC distribution in an SLB containing 5% total PI. The distribution of PC was tracked using TopFluor TMR PC. ATP was added before image acquisition. Initial lipid composition: 94.9% DOPC, 5% 18:1 PI, and 0.1% TopFluor TMR PC. Proteins: 20 nM MavQ, doped with 25% mEGFP-MavQ, and 2 µM SidP. Scale bar: 50 µm.

**Movie S15.**

Time-lapse confocal imaging of MavQ patterns and the underlying distribution of a non-substrate fluorophore-labeled PE in an SLB containing 5% total PI. ATP was added before image acquisition. Initial lipid composition: 95% DOPC, 5% 18:1 PI, and an extra 0.01% ATTO 647N PE. Proteins: 20 nM MavQ, doped with 25% mEGFP-MavQ, and 2 µM SidP. Scale bar: 50 µm.

**Movie S16.**

Time-lapse confocal imaging of MavQ patterns and the underlying PI distribution in an SLB containing 1% total PI. The distribution of PI was tracked using TopFluor TMR PI. ATP was added before image acquisition. Initial lipid composition: 99% DOPC, 0.9% 18:1 PI, and 0.1% TopFluor TMR PI. Proteins: 20 nM MavQ, doped with 25% mEGFP-MavQ, and 2 µM SidP. Scale bar: 50 µm.

**Movie S17.**

Time-lapse confocal imaging of MavQ patterns and the underlying lipid substrate distribution in an SLB containing 1% total PI. The distribution of the combined PI and PI3P pool was tracked using TopFluor TMR PI, while PI3P was specifically visualized using mTagBFP2-2xFYVE_Hrs_. ATP was added before image acquisition. Initial lipid composition: 99% DOPC, 0.9% 18:1 PI, and 0.1% TopFluor TMR PI. Proteins: 20 nM MavQ, doped with 25% mEGFP-MavQ, 2 µM SidP, and 20 nM mTagBFP2-2xFYVE_Hrs_. Scale bar: 50 µm.

**Movie S18.**

Time-lapse confocal imaging of MavQ patterns and the underlying PC distribution in an SLB containing 1% total PI. The distribution of PC was tracked using TopFluor TMR PC. ATP was added before image acquisition. Initial lipid composition: 98.9% DOPC, 1% 18:1 PI, and 0.1% TopFluor TMR PC. Proteins: 20 nM MavQ, doped with 25% mEGFP-MavQ, and 2 µM SidP. Scale bar: 50 µm.

**Movie S19.**

Time-lapse confocal imaging of MavQ patterns and the underlying distribution of a non-substrate fluorophore-labeled PE in an SLB containing 1% total PI. ATP was added before image acquisition. Initial lipid composition: 99% DOPC, 1% 18:1 PI, and an extra 0.01% ATTO 647N PE. Proteins: 20 nM MavQ, doped with 25% mEGFP-MavQ, and 2 µM SidP. Scale bar: 50 µm.

**Movie S20.**

MavQ pattern dynamics from numerical simulations of the reaction–diffusion model at varying bulk MavQ concentration (*M*) and SidP-to-MavQ ratios (*P*/*M*). The average PI + PI3P lipid area fraction ⟨*ϕ*⟩ = 0.15.

**Movie S21.**

MavQ pattern dynamics from numerical simulations of the reaction–diffusion model at varying bulk MavQ concentration (*M*) and average PI + PI3P lipid area fractions (⟨*ϕ*⟩). The bulk SidP- to-MavQ ratio is held at *P*/*M* = 32.

**Movie S22.**

Time-lapse confocal imaging of MavQ pattern formation on a rectangular membrane patch showing wave reflection at the boundaries. ATP was added before image acquisition. Initial lipid composition: 95% DOPC, 5% 18:1 PI, and an extra 0.01% ATTO 647N PE. Proteins: 20 nM MavQ, doped with 25% mEGFP-MavQ, and 1 µM SidP. Scale bar: 50 µm.

**Movie S23.**

Time-lapse confocal imaging of MavQ self-organization on small square corrals and narrow lanes, showing that no coherent MavQ patterns emerge, but MavQ waves rotated within individual corrals and propagated only along the lanes. Initial lipid composition: 95% DOPC, 5% 18:1 PI, and an extra 0.01% ATTO 647N PE. Proteins: 50 nM MavQ, doped with 25% mEGFP-MavQ, and 2 µM SidP. Scale bar: 50 µm.

